# ANP32A represses Wnt signaling across tissues tissues thereby protecting against joint and heart disease

**DOI:** 10.1101/2021.04.04.438364

**Authors:** Silvia Monteagudo, Frederique M. F. Cornelis, Xiangdong Wang, Astrid de Roover, Tine Peeters, Jolien Quintiens, An Sermon, Rodrigo C. de Almeida, Ingrid Meulenbelt, Rik J. Lories

**Affiliations:** Laboratory of Tissue Homeostasis and Disease, Skeletal Biology and Engineering Research Center, Department of Development and Regeneration, KU Leuven, Leuven, Belgium; Department of Trauma Surgery, University Hospitals Leuven, Leuven, Belgium; Department of Development and Regeneration, KU Leuven, Leuven, Belgium; Department of Biomedical Data Sciences, Section of Molecular Epidemiology, Leiden University Medical Center, RC Leiden, The Netherlands; Integrated research on Developmental determinants of Ageing and Longevity (IDEAL), RC Leiden, The Netherlands; Division of Rheumatology, University Hospitals Leuven, Leuven, Belgium

**Keywords:** Cardiac hypertrophy/ Cartilage / Osteoarthritis/ Rheumatology/ Wnt signaling

## Abstract

Wnt signaling is key to diverse homeostatic and pathological processes. This cascade is hyper-activated in osteoarthritis, the most common joint disease. Yet, fundamental aspects of Wnt signaling remain undiscovered. Here, we report that ANP32A negatively regulates Wnt signaling across tissues. In cartilage, loss of *Anp32a* triggered Wnt hyper-activation. Mechanistically, ANP32A directly interacted with Wnt pathway components and inhibited Wnt target genes via histone acetylation masking. Wnt antagonist treatment reduced severity of osteoarthritis in *Anp32a*-deficient mice preventing osteophyte formation, contrasting with cartilage-protective effects of ANP32A on oxidative stress. Hence, dual therapy targeting Wnt signaling and oxidative stress in *Anp32a*-deficient mice ameliorated more osteoarthritis features than individual treatments. *Anp32a* loss also resulted in Wnt hyper-activation in the heart with cardiac hypertrophy, and in the hippocampus, shedding light on mechanisms for reported links between ANP32A and Alzheimer’s disease. Collectively, this work reveals that ANP32A is a translationally relevant repressor of Wnt signaling, impacting homeostasis and disease across tissues.

## Introduction

Tissue growth and homeostasis are processes that require an orchestrated activation and restriction of gene expression programs. How specific genes are switched on and off at the correct time and in the right place is a central question in biology. Furthermore, understanding how gene regulatory processes might be perturbed by disease or in the presence of biological factors such as genetic variants is key to establish the rationale for the development of effective disease-modifying treatments and personalized medicine.

We previously discovered that the acidic leucine rich nuclear phosphoprotein-32A (*ANP32A*) protein switches on the transcription of the ataxia telangiectasia mutated serine threonine kinase (*ATM*) gene to tightly control a central regulatory network that prevents oxidative processes in cartilage, cerebellum and bone (Cornelis et al., 2018). In cartilage, a specialized semi-rigid and flexible connective tissue that covers the ends of the bones within the joints and that is critical for normal mobility, absence of ANP32A leads to severe osteoarthritis and oxidative stress (Cornelis et al., 2018). Earlier, genetic variants in *ANP32A* were associated with this disease (Valdes et al., 2009; Valdes et al., 2008). Osteoarthritis is the most common age-related or trauma-triggered chronic joint disorder worldwide (Hunter et al., 2014), and a leading source of enduring pain and disability (Murray et al., 2013; Vos et al., 2012). It is primarily characterized by cartilage destruction, as well as by osteophyte formation, synovial inflammation, and abnormal subchondral bone remodeling. Current treatments for osteoarthritis are limited to symptom relief and in advanced cases joint replacement surgery may be the only option. Thus, developing an effective therapy that arrests or reverses disease progression is urgently needed. As osteoarthritis is a disease of the whole joint affecting different tissues, distinct pathways with varying downstream pathological effects are likely involved and effective treatment may require combination therapies.

Importantly, absence of ANP32A not only leads to osteoarthritis, but also to cerebellar ataxia and osteopenia (Cornelis et al., 2018). These discoveries revealed that ANP32A is a pleiotropic protein, influencing several phenotypic traits. Pleiotropic proteins are central in protein-protein interaction networks and may control multiple biological pathways (Ittisoponpisan et al., 2017). Thus, key unanswered questions are whether ANP32A has a role in multiple molecular mechanisms of homeostasis and disease, and whether there are yet undiscovered phenotypic traits critically influenced by ANP32A.

Here, we identify that ANP32A adjusts the transcriptional response of Wnt signaling, a central network in tissue homeostasis and disease. We provide biochemical evidence unravelling that ANP32A performs this regulatory role via a histone acetylation masking mechanism. From a translational perspective, we evaluate the disease-modifying effects of targeting Wnt hyper-activation in a model of osteoarthritis in *Anp32a*-deficient mice, and the clinical benefits of a combination with antioxidant treatment in this setting. Additionally, we explore whether ANP32A regulates Wnt signaling in tissues beyond cartilage. Our findings unveil that ANP32A is a major regulatory molecule of at least two central biological pathways across multiple tissues, with new links to osteoarthritis and other pathologies, namely cardiac hypertrophy and Alzheimer’s disease.

## Results

### Transcriptome analysis of articular cartilage suggests dysregulation of Wnt signaling in *Anp32a*-deficient mice

To explore whether ANP32A regulates central networks in cartilage biology beyond oxidative stress, we sought to identify the signaling cascades that drive differences in gene expression in articular cartilage from *Anp32a*^-/-^ mice (mice with a global deletion of ANP32A) compared to wild-type mice. We took advantage of our earlier microarray transcriptome study (geonr: GSE108036) (Cornelis et al., 2018), and applied the Upstream Regulator Analysis approach from Ingenuity Pathway Analysis (IPA) software. Pathway analysis of identified genes classified as “transcriptional regulators” indicated that p53 signaling, which is directly linked to ATM, is the main enriched pathway mediating loss of ANP32A’s effects on gene expression programs (Figure 1A), corroborating our earlier observations (Cornelis et al., 2018). Strikingly, the analysis also suggested that the Wnt signaling pathway drives gene expression changes resulting from loss of ANP32A (Figure 1A).

**Figure 1.**
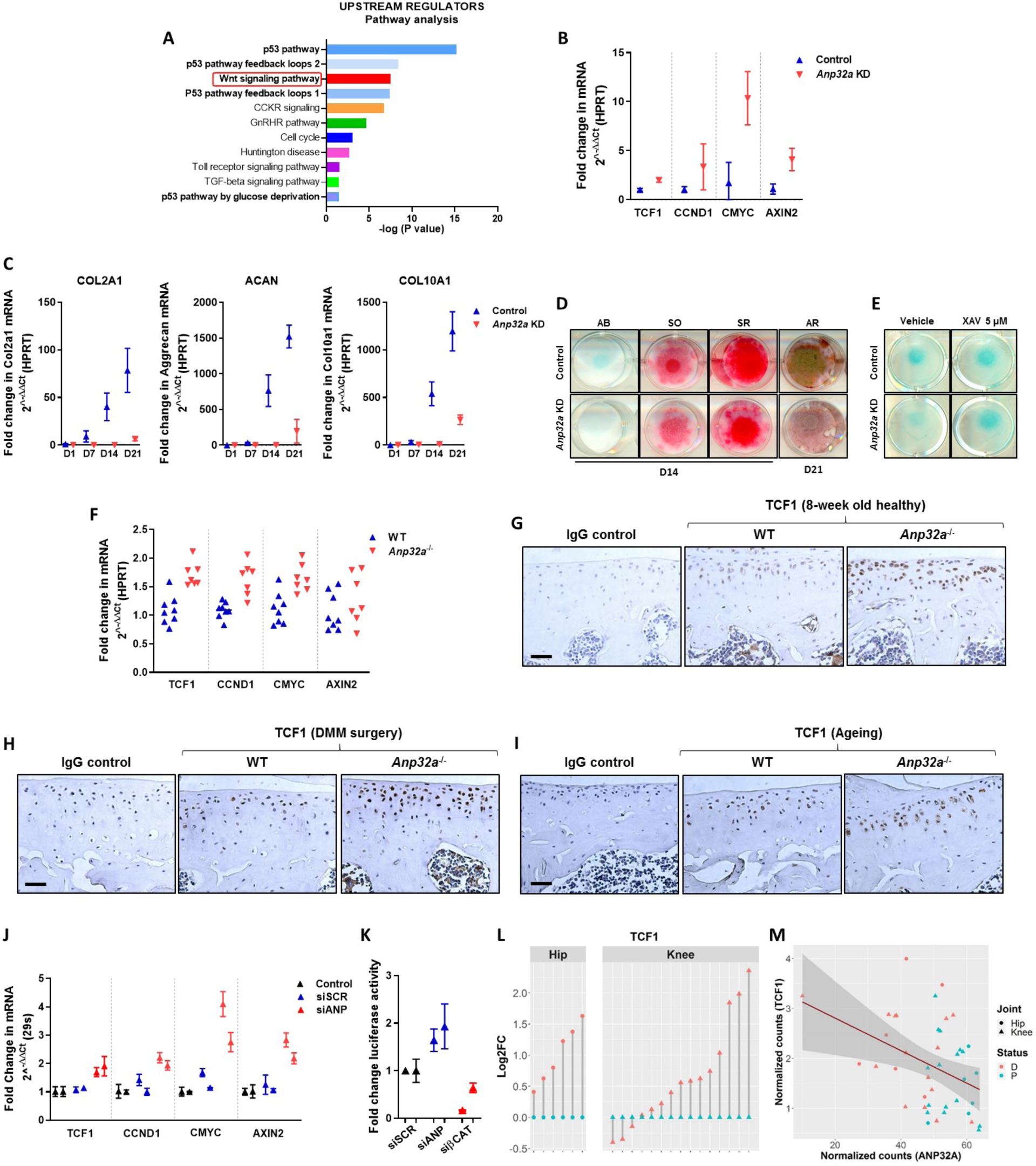
Loss of ANP32A triggers Wnt signaling hyper-activation in cartilage. **(A)** PANTHER pathway analysis of upstream transcriptional regulators identified in microarray data comparing articular cartilage of 8-week old male *Anp32a*-deficient to wild-type mice (n = 4 per group) using Ingenuity Pathway Analysis (IPA). **(B)** Real-time PCR analysis of direct Wnt target genes *Tcf1*, *Ccnd1*, *cMyc* and *Axin2* in control and *Anp32a* knockdown (KD) ATDC5 cells (Pillai = 0.964, F_2,3_ = 40.34, *P* = 0.0068 by MANOVA; *Ccnd1* and *Axin2* showed ≥ 0.9 correlation with *Tcf1* and were not included in the model, mean ± SD of 3 technical replicates per condition). **(C)** Real-time PCR analysis of chondrogenic differentiation markers collagen 2 (*Col2a1*), aggrecan (*Acan*) and collagen 10 (*Col10a1*) in control and *Anp32a* KD ATDC5 cells [F_3,12_ = 25.321, *P* < 0.0001 (*Col2a1*), F_1.05,4.21_ = 24.378 *P* = 0.007 (*Acan*), F_3,12_ = 42.775 *P* < 0.0001 (*Col10a1*) by 2-way ANOVA for interaction between silencing and time, mean ± SD of 3 technical replicates per condition]. **(D)** Alcian blue (AB), safranin O (SO), picrosirius red (SR) at day 14 (D14) and alizarin red (AR) staining at day 21 (D21) showing reduced proteoglycan deposition (AB, SO), collagen content (SR) and mineralization (AR) in *Anp32a* KD ATDC5 cells during chondrogenesis. **(E)** AB staining demonstrating rescue of chondrogenic differentiation in *Anp32a* KD ATDC5 cells by treatment with Wnt inhibitor XAV939 (XAV). **(F)** Real-time PCR analysis of Wnt target gene expression in articular cartilage from 8-week old male wild-type (WT) and *Anp32a*-deficient (*Anp32a^-/-^*) mice (Pillai = 0.788, F_3,11_ = 13.61, *P* = 0.0005 by MANOVA – *cMyc* showed ≥ 0.9 correlation with *Tcf1* and was not included in the model, n = 8 and 7 mice per group). **(G-I)** Immunohistochemical staining for TCF1 protein in male 8-week old (G), osteoarthritic (H) and female ageing WT and *Anp32a^-/-^* mice (I) (representative images of n = 3 different mice per group). Scale bar 50 µm. **(J)** Real-time PCR analysis of Wnt target gene expression in human articular chondrocytes transfected with siRNA targeting *ANP32A* (siANP32A) or scrambled siRNA (siSCR) (n = 2 different donors - mean ± SD of 3 technical replicates). **(K)** TOP/FOP reporter assay in human articular chondrocytes transfected with siANP32A, siRNA targeting β-catenin (siβCAT) or siSCR, and then treated with recombinant WNT3A (n = 2 biologically independent experiments, mean ± SD of 3 technical replicates). **(L-M)** *TCF1* expression (L) and negative correlation with *ANP32A* expression (M) by RNA sequencing in paired preserved and damaged cartilage from hips (o) and knees (Δ) from osteoarthritis patients [log2-fold change (Log2FC) of damaged (D) *vs.* preserved (P)) (n = 21, *P* < 0.0001, Benjamini-Hochberg adjusted paired *t*-test (l), Pearson’s correlation = -0.40 - *P* = 0.0084 (m)].

### *Anp32a* deficiency impairs chondrogenic differentiation via Wnt hyper-activation

Combined genetic and experimental evidence strongly supports that a fine-tuned balance of Wnt activity is key for cartilage health, and that excessive activation of Wnt signaling is harmful and contributes to osteoarthritis (Monteagudo & Lories, 2017; Zhu et al., 2009). To explore whether ANP32A has a regulatory effect on Wnt signaling in cartilage, we first used an *in vitro* cartilage differentiation model in which Wnt signaling plays a key role. The chondrogenic ATDC5 cell line exhibits a multistep differentiation process towards cartilage when the cells are seeded in micromasses (Yao & Wang, 2013). In this model, Wnt signaling activation in the early phase blocks the differentiation program (Wang et al., 2019). We generated stable *Anp32a* siRNA knockdown (KD) ATDC5 cell lines (Supplementary Figure 1A) and found that gene expression levels of direct Wnt target genes were up-regulated in these *Anp32a*-deficient cells compared to controls (Figure 1B). Upon micromass differentiation culture, mRNA expression of early cartilage differentiation markers *collagen 2* (*Col2a1)* and *aggrecan (Acan),* and terminal differentiation marker *collagen 10* (*Col10a1)* were strongly down-regulated in *Anp32a* KD cells compared to controls (Figure 1C). To evaluate proteoglycan and collagen content of the micromasses, we performed alcian blue, safranin O and picrosirius red staining. We observed a reduction in proteoglycans and collagen amounts in *Anp32a* KD micromasses compared with controls (Figure 1D). Mineralization of the micromasses, assessed by alizarin red staining, was also reduced (Figure 1D). Although expression of *Atm* was partially suppressed in *Anp32a* KD micromasses compared with controls (Supplementary Figure 1B), treatment of *Anp32a* KD micromasses with antioxidant N-acetylcysteine (NAC) did not rescue chondrogenic marker expression (Supplementary Figure 2A). However, blockade of β-catenin dependent Wnt signaling with XAV939 (XAV) (Huang et al., 2009) showed rescue effects on chondrogenic differentiation in *Anp32a* KD cells (Figure 1E, Supplementary Figure 2B). These observations indicate that active Wnt signaling contributes to the detrimental effects of loss of *Anp32a* in this cartilage differentiation model.

### *Anp32a* deficiency leads to hyper-activation of Wnt signaling in mouse and human articular cartilage

We then investigated whether Wnt signaling is similarly dysregulated in articular cartilage of *Anp32a*^-/-^ mice, both at the gene and protein levels. We found that mRNA amounts of direct Wnt target genes were up-regulated in *Anp32a*^-/-^ mice compared to wild-types (Figure 1F). Furthermore, protein reactivity of Wnt target gene TCF1 was also increased in articular cartilage from *Anp32a*^-/-^ mice, as assessed by immunohistochemistry, in healthy young mice at 8 weeks (Figure 1G), after induction of osteoarthritis using the destabilization of the medial meniscus model (DMM) (Figure 1H) and upon ageing (12 months) (Figure 1I). In human articular chondrocytes, expression of Wnt target genes was also up-regulated upon siRNA-mediated *Anp32a* silencing (Figure 1J). In addition, *Anp32a*-silenced human articular chondrocytes showed stronger induction of the Wnt/β-catenin pathway reporter TOP-FLASH than control cells, upon recombinant WNT3A stimulation (Figure 1K). All these findings indicate that ANP32A negatively regulates Wnt signaling in articular cartilage.

Of note, in patients with knee and hip osteoarthritis, *ANP32A* expression was down-regulated in damaged as compared to preserved areas of the articular cartilage (Cornelis et al., 2018). Conversely, expression of Wnt direct target genes was up-regulated in damaged areas of osteoarthritis cartilage (Figure 1L and Supplementary Figure 3A). Expression of *ANP32A* and several Wnt target genes negatively correlated in human osteoarthritis cartilage (Figure 1M and Supplementary Figure 3B). These data indicate that the link between ANP32A and Wnt signaling may be clinically relevant in the pathogenesis of osteoarthritis.

### Active Wnt signaling triggers nuclear translocation of ANP32A

We next sought to investigate the molecular mechanism via which ANP32A negatively regulates Wnt signaling. The core of the canonical or Wnt/ β-catenin cascade is the regulation of β-catenin protein levels by a cytoplasmic destruction complex. In the absence of Wnt signaling, the destruction complex captures cytosolic β-catenin, leading to its phosphorylation by GSK3 and subsequent degradation by the proteasome. Wnt receptor activation leads to functional inactivation of the destruction complex, resulting in β-catenin accumulation and nuclear entry, where it binds to TCF/LEF transcription factors and regulates the transcription of Wnt target genes (Clevers, 2006). A high throughput yeast two-hybrid screening demonstrated interactions between ANP32A and AXIN1, a central scaffold for the components of the destruction complex (Stelzl et al., 2005). We thus investigated whether ANP32A interacts with AXIN1 at the protein level in articular chondrocytes. We carried out co-immunoprecipitations of endogenous ANP32A from extracts of cells treated with recombinant WNT3A ligand or vehicle control. This analysis revealed that ANP32A bound AXIN1 in baseline conditions, and this association was abolished upon Wnt signaling activation by WNT3A (Figure 2A).

**Figure 2.**
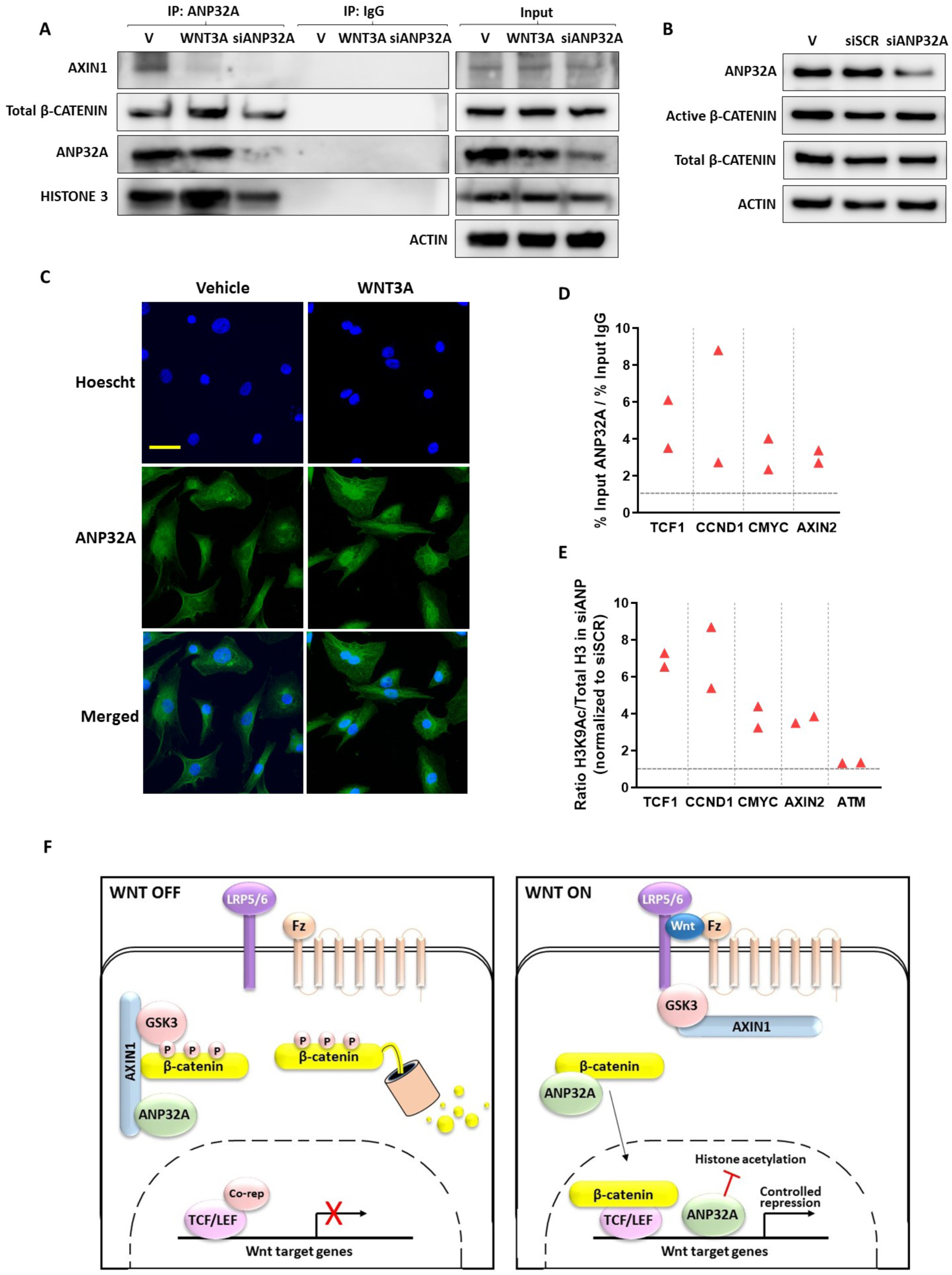
ANP32A represses Wnt target gene expression by histone acetylation masking. **(A)** Co-immunoprecipitation (Co-IP) using an anti-ANP32A antibody showing the binding between ANP32A and AXIN1 in untreated human articular chondrocytes, which is abolished upon Wnt activation by recombinant WNT3A protein. Wnt activation increases the binding of ANP32A with β-catenin and with histone 3. Silencing of ANP32A (siANP32A) is shown to assess the specificity of the ANP32A antibody used in the immunoprecipitation. The image is representative of three independent experiments. **(B)** Immunoblot analysis of ANP32A, active β-catenin and total β-catenin protein amounts (with actin as loading control) in human articular chondrocytes transfected with *ANP32A* (siANP32A) or scrambled siRNA (siSCR). **(C)** Immunofluorescent staining of ANP32A (green) and counterstaining of nuclei with Hoechst dye (blue) in human articular chondrocytes upon vehicle or recombinant WNT3A treatment (4 hours). Representative images are shown (n = 2). Scale bar 10 µm. **(D)** Chromatin-immunoprecipitation quantitative PCR (ChIP-qPCR) analysis of ANP32A binding to chromatin on Wnt target (*Tcf1*, *Cnnd1,cMyc* and *Axin2*) gene promoter regions in human articular chondrocytes with high endogenous Wnt signaling (cells were expanded two passages). **(E)** ChIP-qPCR analysis of acetylated H3K9 (H3K9Ac) on Wnt target (*Tcf1*, *Ccnd1, cMyc* and *Axin2*) and *Atm* gene promoter regions in human articular chondrocytes with high endogenous Wnt signaling that were transfected with siANP32A or siSCR. Data are expressed as ratio of H3K9Ac and total Histone 3 (H3) in siANP32A cells normalized to siSCR cells. Data points are from two biologically independent experiments (D-E). **(F)** Scheme summarizing how ANP32A regulates the Wnt transcriptional response. In basal conditions (left panel), ANP32A interacts with AXIN1 in the destruction complex. Upon Wnt activation (right panel), ANP32A dissociates from AXIN1, associates with β-catenin, and translocates to the nucleus. Within the nucleus, ANP32A represses Wnt target genes via blocking histone acetylation.

These observations prompted us to investigate the functional relevance of the interaction of ANP32A with AXIN1 for the cytoplasmic stabilization of β-catenin. Silencing of ANP32A did not affect β-catenin protein levels (active nor total) in human articular chondrocytes (Figure 2B). These data suggest that ANP32A may regulate Wnt signaling downstream of β-catenin stabilization. Remarkably, the dissociation of ANP32A from AXIN1 upon Wnt activation paralleled an enhanced association of ANP32A with β-catenin (Figure 2A). Then, we investigated whether Wnt activation triggers nuclear translocation of ANP32A, as it occurs for β-catenin. We examined the effect of recombinant WNT3A ligand on the subcellular localization of endogenous ANP32A by immunofluorescence in human articular chondrocytes. ANP32A was mostly cytoplasmic in untreated cells (Figure 2C), in agreement with our previous data (Cornelis et al., 2018). In contrast, ANP32A was also markedly localized in the nucleus after WNT3A treatment (Figure 2C).

### ANP32A represses Wnt target gene expression through histone acetylation masking

Next, we sought to investigate how ANP32A represses Wnt target genes. ANP32A is a member of the inhibitor of histone acetyltransferase (INHAT) complex (Seo et al., 2001), which limits transcription by binding to histones, preferentially histone 3, and sterically hindering acetylation (Reilly et al., 2014). Hypoacetylation of histones is linked to condensed chromatin and transcriptional repression (Verdone et al., 2005). We found that Wnt-induced ANP32A nuclear accumulation in human articular chondrocytes (Figure 2C) was paralleled by an increased interaction of ANP32A with histone 3 (Figure 2A), indicating that ANP32A may repress Wnt target gene expression via its inhibitory role on histone acetylation. Chromatin-immunoprecipitation (ChIP)-qPCR analysis showed that ANP32A bound the chromatin at Wnt target gene promoters in human articular chondrocytes (Figure 2D). In these cells, silencing of ANP32A resulted in increased H3K9 acetylation at Wnt target gene promoters (Figure 2E). Collectively, these results indicate that Wnt induces ANP32A nuclear internalization and chromatin binding to Wnt target gene promoters, resulting in inhibition of histone acetylation, and thus, transcriptional repression of Wnt target genes (see scheme in Figure 2F). Of note, in *Anp32a*-silenced cells, we did not find increased H3K9 acetylation marks at the promoter of *Atm* (Figure 2E), a gene that is positively regulated at the transcriptional level by ANP32A (Cornelis et al., 2018).

### Targeting Wnt hyper-activation and oxidative stress downstream of ANP32A deficiency affects different features of osteoarthritis

Next, we investigated the therapeutic implications of our findings for osteoarthritis. *Anp32a* deficiency results in hyper-activation of Wnt signaling in articular cartilage (Figure 1G), and this occurs concomitantly with an increase in oxidative stress (Cornelis et al., 2018). We evaluated the effects of treatment with a Wnt inhibitor and an antioxidant, individually or in combination, on osteoarthritis, using the DMM mouse model of the disease (Glasson et al., 2007) that mimics mechanisms and features of posttraumatic osteoarthritis in humans (Lotz & Kraus, 2010). In this model, we previously showed that *Anp32a*^-/-^ mice have more severe cartilage damage and oxidative stress compared to wild-type controls and sham-operated *Anp32a*^-/-^ mice (Cornelis et al., 2018). NAC treatment partially prevented the increased severity of osteoarthritis compared to wild-types after induction of the model (Cornelis et al., 2018). In a new experiment, we demonstrate Wnt hyper-activation in the articular cartilage of DMM-operated Anp32a-/- mice (Figure 1H). After DMM surgery, *Anp32a*^-/-^ mice were now intra-articularly injected with Wnt inhibitor XAV and were given NAC via the drinking water as before (Figure 3A). XAV treatment did not provide cartilage protection (Figures 3B-C), although it effectively inhibited Wnt hyper-activation in articular cartilage (Figure 3D). However, we found that Wnt inhibition prevented osteophyte formation in the osteoarthritis model in *Anp32a*^-/-^ mice (Figure 3E-G). In contrast, NAC protected the *Anp32a*^-/-^ mice against cartilage damage but had no effect on osteophyte formation (Figures 3B-G). The combination of XAV and NAC improved both cartilage damage and osteophyte formation (Figures 3B-G). These observations suggest that the pathological consequences of ANP32A deficiency may hinge on different pathways depending on the tissue involved, in this case oxidative stress in articular cartilage and hyper-activation of Wnt signaling in osteophyte formation.

**Figure 3.**
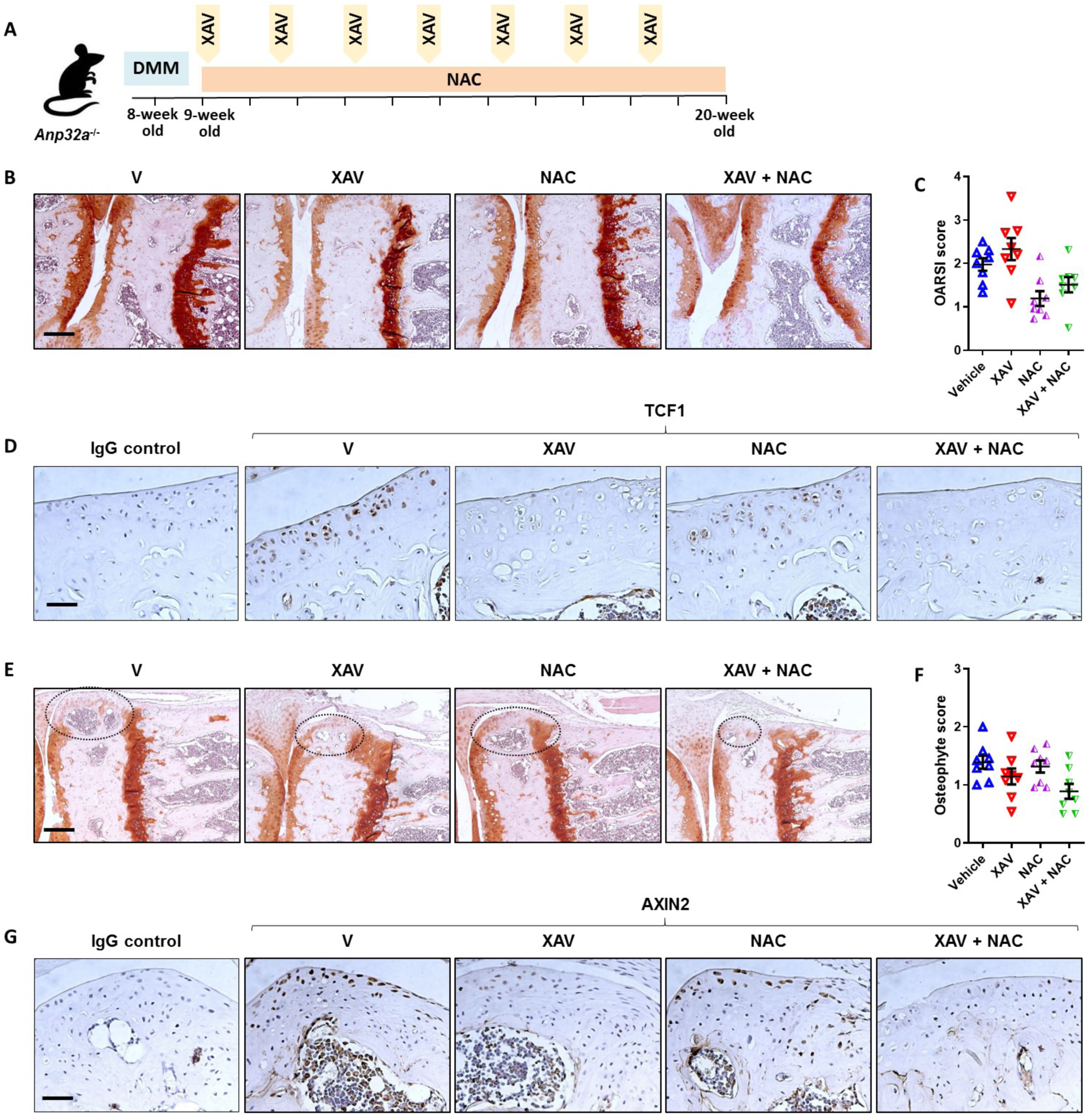
ANP32A protects against different features of osteoarthritis by controlling distinct pathways. **(A)** Schematic outline of *in vivo* pharmacological interventions against osteoarthritis in *Anp32a*-deficient (*Anp32a^-/-^*) male mice. 8-week old mice were subjected to destabilization of the medial meniscus (DMM) surgery. One week after injury, mice were injected intra-articularly with vehicle or Wnt inhibitor XAV939 (XAV) every 10 days for a total of 7 times. Mice were treated orally with vehicle, XAV or NAC alone or in combination. Knee joints were collected 12 weeks after surgery. **(B-C)** Hematoxylin-safranin-O-stained sections (B) and quantification by OARSI severity grade (C) demonstrating that NAC protects against articular cartilage damage in osteoarthritis: [F_1,28_=8.48 *P* = 0.007 for main effect NAC by two-way ANOVA, n = 8 per group]. Scale bar 200 µm. **(D)** Immunohistochemical staining for TCF1 protein in articular cartilage of *Anp32a^-/-^* mice in the DMM model treated or not with XAV or NAC (representative images of n = 5 different mice per group). Scale bar 50 µm. (**E-F)** Hematoxylin-safranin-O-stained sections (E) and quantification (F) of osteophytes demonstrating that Wnt inhibition protects against osteophyte formation in osteoarthritis [F_1,28_=7.69 *P* = 0.0098 for main effect XAV by two-way ANOVA, n = 8 per group]. Scale bar 200 µm. **(G)** Immunohistochemical staining for TCF1 protein in developing osteophytes in *Anp32a^-/-^* mice in the DMM model treated or not with XAV or NAC (representative images of n = 5 different mice per group). Scale bar 50 µm.

### ANP32A negatively regulates Wnt signaling in heart and hippocampus

As demonstrated above, ANP32A interacts with different components of the Wnt signaling pathway. Highly-connected nodes within a signaling pathway, which moreover are ubiquitously expressed like ANP32A (Uhlén et al., 2015; Wang et al., 2015) (Human Protein Atlas available from http://www.proteinatlas.org), may regulate such cascade in multiple tissues (Freeman et al., 2015). Thus, we next aimed to investigate whether ANP32A’s regulatory role on Wnt signaling is relevant beyond the joint. We analyzed data from the Human Protein Atlas to identify tissues and organs with an inverse relationship between the expression of ANP32A and Wnt target gene TCF1, which may indicate an underlying potential regulatory link between ANP32A and Wnt signaling. Although there was no ubiquitous relationship between these factors, our analysis identified brain and muscle including the heart, as organs of interest (Figure 4A).

**Figure 4.**
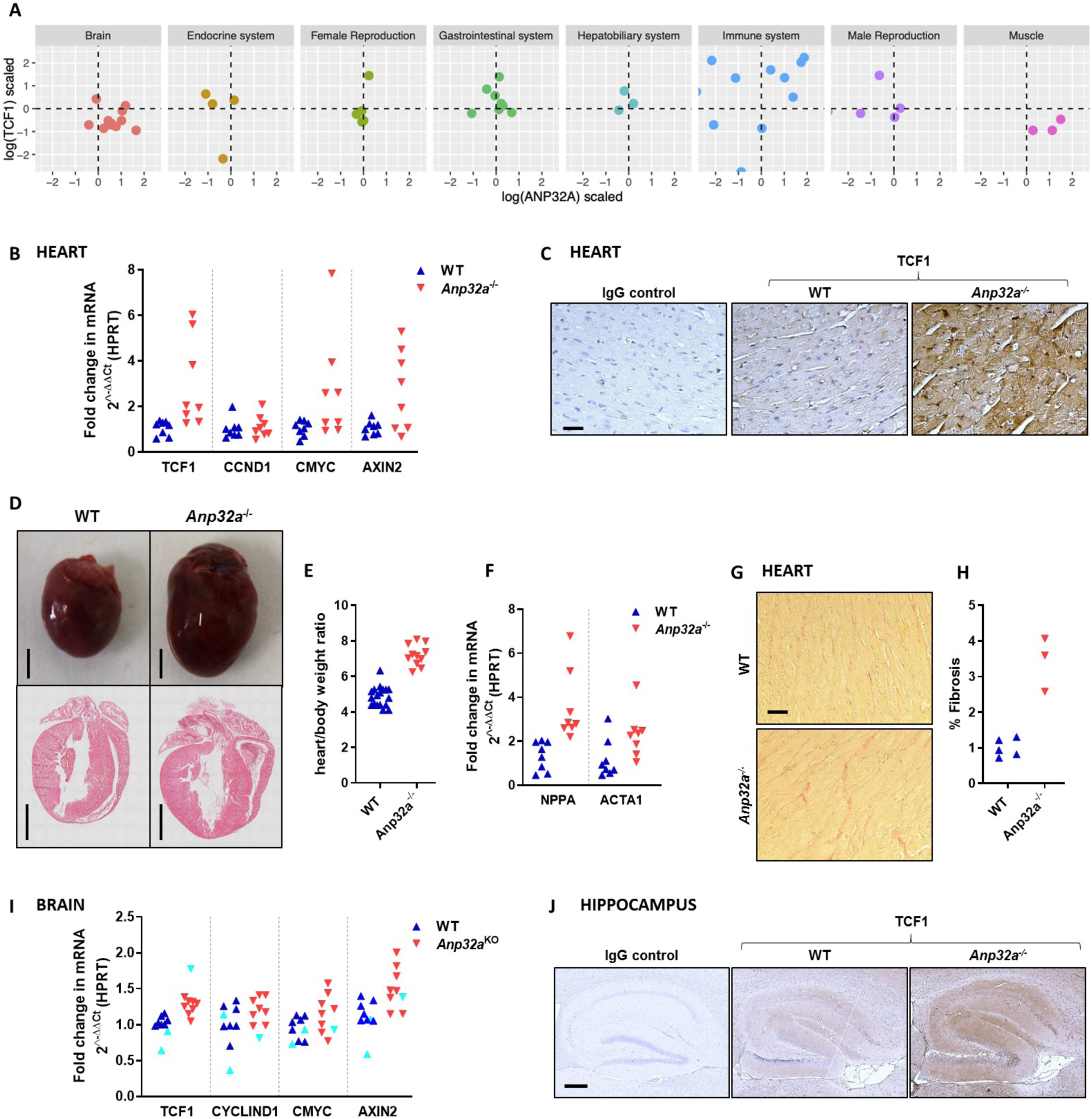
Regulation of Wnt signaling by ANP32A in tissues beyond cartilage. **(A)** Selected Protein Atlas data showing the inverse relationship between ANP32A and Wnt target gene TCF1 in brain and muscle tissues compared to other systems. Data are presented as scaled variables. **(B)** Real-time PCR analysis of Wnt target gene expression (n = 8) in heart from 20-week old male WT and *Anp32a^-/-^* mice (Pillai = 0.492, F_3,12_ = 3.877, *P*=0.038 by MANOVA - *Axin2* data did not show homogeneity of variance and were not included in the model). **(C)** Immunohistochemical staining for TCF1 in 20-week old male WT and *Anp32a^-/-^* mice hearts. Scale bar 50 µm. **(D)** Macroscopic and Hematoxylin-Eosin stained hearts demonstrating cardiac hypertrophy in *Anp32a^-/-^* compared to WT mice. Scale bar 2 mm. (**E**) Heart weight/body weight ratio in 20-week old male WT and *Anp32a^-/-^* mice (t_27_ = 10.45, *P <* 0.0001). (**F**) Real-time PCR analysis of hypertrophy markers *natriuretic peptide precursor A (Nppa)* and *skeletal muscle α- actin (Acta1)* (n = 8) in heart from 20-week old male WT and *Anp32a^-/-^* mice (t_14_ = 4.010, *P* = 0.0011 (*Nppa*), t_14_ = 2.87, *P =* 0.0125 (*Acta1*) by Student’s t-test). (**G-H**) Picrosirius red staining (G) showing increased amounts of fibrotic tissue in hearts from 20-week old *Anp32a^-/-^* mice compared to WT (H) (t_6_ = 6.82, *P* = 0.0005 by Student’s t-test). Scale bar 50 µm. **(I)** Real-time PCR analysis of Wnt target gene expression (n = 9) in brain from 8-week old male WT and *Anp32a^-/-^* mice (Pillai = 0.666, F_4,10_ = 4.978, *P*=0.018 by MANOVA – 3 extreme outliers were excluded from the model as indicated in cyan color). **(J)** Immunohistochemical staining for TCF1 in 16-week old male hippocampus from WT and *Anp32a^-/-^* mice. Scale bar 250 µm.

In heart tissue, Wnt signaling was enhanced in *Anp32a*^-/-^ mice compared to controls, both at the gene and the protein expression levels (Figures 4B-C). Expression of *Anp32a* was shown to be down-regulated in a study exploring key deregulated genes and pathways involved in cardiac hypertrophy (Gao et al., 2018). Wnt signaling is known to be activated during heart failure and cardiac hypertrophy (Bergmann, 2010; Stylianidis et al., 2017). We found that hearts from 20-week old male *Anp32a*^-/-^ mice were enlarged compared to controls (Figure 4D). In *Anp32a*^-/-^ animals, total heart weight as well as heart weight expressed relative to body weight were higher than in wild-type mice (Figure 4E and Supplementary Table 1). The expression of hypertrophy markers natriuretic peptide precursor A *(Nppa)* and skeletal muscle α-actin *(Acta1)* was up-regulated in *Anp32a*^-/-^ compared to control mice (Figure 4F) and histology analysis showed increased amounts of fibrotic tissue (Figures 4G-H). Collectively, our data show that lack of ANP32A triggers the development of spontaneous cardiac hypertrophy, which may result from an excessive activation of Wnt signaling.

As suggested by the Protein Atlas data analysis, Wnt target gene expression was up-regulated in brain from *Anp32a*^-/-^ mice compared to controls, demonstrating that ANP32A negatively regulates Wnt signaling in this organ (Figure 4I). Importantly, ANP32A dysregulation has been linked to Alzheimer’s disease. ANP32A is increased in human brains from Alzheimer’s disease patients (Tanimukai et al., 2005; Tsujio et al., 2005). In a mouse model of this disease, ANP32A elevation in the hippocampus correlates with learning deficits, and downregulating ANP32A rescues synaptic plasticity and memory loss (Chai et al., 2017; Feng et al., 2017). Conversely, overexpression of ANP32A in hippocampus induced memory deficits in mice (Chai et al., 2018). Loss of Wnt signaling plays a critical role in the pathogenesis of Alzheimer’s disease and emerging studies suggest that restoring Wnt signaling may be a promising therapeutic strategy (De Ferrari et al., 2014; Jia et al., 2019; Tapia-Rojas & Inestrosa, 2018). Remarkably, we found that Wnt signaling was enhanced in the hippocampus of *Anp32a*^-/-^ mice (Figure 4J). Therefore, our insights suggest that ANP32A’s involvement in the pathogenesis of Alzheimer’s disease may be linked to a decline in activation of Wnt signaling.

## Discussion

Our findings reveal that ANP32A is a converging node regulating the transcriptional responses of two central cascades in cell biology: the Wnt signaling pathway as identified here, and oxidative stress as we previously showed (Cornelis et al., 2018). Thus, ANP32A is proposed as a key regulator of a complex network of pathways that protect against osteoarthritis. We also provide evidence that the regulatory role of ANP32A on the Wnt transcriptional response is not restricted to cartilage, but also functions in tissues where dysregulation of *ANP32A* expression has been linked to disease, namely heart and hippocampus.

In the absence of ANP32A, Wnt signaling is hyper-activated in a cartilage differentiation model, and in mouse and human articular cartilage. Expression of *ANP32A* and several Wnt target genes negatively correlates in human articular cartilage from osteoarthritis patients, indicating that dysregulation of the ANP32A-Wnt axis may play a key pathophysiological role in osteoarthritis development and progression. The Wnt/β-catenin pathway has been extensively implicated in the pathogenesis of osteoarthritis (Corr, 2008; Lories & Monteagudo, 2020; Monteagudo & Lories, 2017). In humans, polymorphisms in genes involved in Wnt signaling, particularly in the genes encoding extracellular inhibitor *SFRP3* and epigenetic modulator *DOT1L,* are associated with increased susceptibility to the development of osteoarthritis (Castaño Betancourt et al., 2012; Loughlin et al., 2004). In rodent models, loss of molecules that suppress Wnt signaling triggers osteoarthritis (Cornelis et al., 2019; Lories et al., 2007; Monteagudo et al., 2017; Nalesso et al., 2017). Furthermore, mechanical injury and inflammation are demonstrated to be potent inducers of Wnt signaling in cartilage (Dell’Accio et al., 2006). All this evidence justifies that Wnt signaling is increasingly recognized as a potential target for osteoarthritis, and clinical trials with Wnt inhibitors are currently underway (Lories & Monteagudo, 2020).

Our preclinical data in the *Anp32a*-deficient genetic mouse model show that combinatorial treatment with a Wnt inhibitor and an antioxidant leads to increased therapeutic efficacy in osteoarthritis. Whereas cartilage damage in the absence of ANP32A is linked to oxidative stress, joint remodeling with osteophyte formation is Wnt dependent. Antioxidants have been successfully used to provide protection for cartilage in translationally relevant pre-clinical settings (Beecher et al., 2007; Nakagawa et al., 2010). Also, in our previous study, NAC treatment showed protective effects for osteoarthritis in *Anp32a*-deficient mice (Cornelis et al., 2018). However, combination of antioxidants and Wnt inhibitors for this disease has never been explored so far. The different effects of the combinatory strategy appear to be translationally relevant for disease management as it demonstrates the potential of combination therapies for osteoarthritis with different interventions simultaneously targeting distinct disease-associated networks and disease manifestations. Given the complexity of osteoarthritis, single target therapies will likely not halt all disease features.

We combined oral antioxidant administration with intra-articular administration of a Wnt inhibitor in mice. The substantial risk of systemic toxicity has prompted a paradigm shift in osteoarthritis drug development with redirection of attention to the benefits of localized versus systemic pharmacological treatment, indicated by the increasing number of intra-articularly administered compounds entering clinical trials (Karsdal et al., 2016). However, locally-administered pharmacological agents in the synovial joint might not easily reach regions closer to the subchondral bone, as dense extracellular matrix in articular cartilage restricts the penetration and diffusion of solutes (DiDomenico et al., 2018). Systemic administration of well tolerated pharmacological agents, such as NAC, might add some benefits to local administration within the joint, as therapeutic effects would not be only dependent on diffusion into the cartilage from the synovial fluid. Once in the blood-stream, small molecules can diffuse from the blood vessels in the subchondral bone into calcified and non-calcified articular cartilage (Arkill & Winlove, 2008; DiDomenico et al., 2018).

Mechanistically, we show that ANP32A regulates Wnt signaling by interacting with chromatin and inhibiting histone acetylation at Wnt target gene promoters, thereby resulting in gene repression. This repressive function is in line with its earlier defined role as part of the INHAT complex, a multiprotein complex that sterically inhibits histone acetyltransferases by binding to histone tails (Schneider et al., 2004; Seo et al., 2002; Seo et al., 2001). Conversely, this molecular mechanism contrasts with our previously reported findings for the role for ANP32A as a positive transcriptional regulator of the *Atm* gene and a previous report indicating that ANP32A enhances gene transcription of interferon-stimulated genes (Cornelis et al., 2018; Kadota & Nagata, 2011). Collectively, these observations indicate that ANP32A can either activate or repress relevant gene expression programs.

We also provide evidence showing that Wnt signaling is enhanced in the heart of *Anp32a*-deficient mice. We examined the heart tissue as Wnt target gene *TCF1* and *ANP32A* expression showed an inverse relationship, and *Anp32a* expression was reported to be downregulated in cardiac hypertrophy (Gao et al., 2018). Besides increased Wnt target gene expression in heart, we also found that loss of *Anp32a* resulted in an increased heart/body weight ratio in mice and more fibrosis. Wnt signaling is activated in cardiac hypertrophy and several studies have reported anti-hypertrophic effects for Wnt inhibitors (Bergmann, 2010). Our findings suggest that ANP32A may protect the heart against the development of cardiac hypertrophy via limiting Wnt signaling. Thus, increasing ANP32A expression may be beneficial in this context.

In this study, we further show that Wnt signaling is enhanced in hippocampus of *Anp32a*-deficient mice. In a genetic mouse model of Alzheimer’s disease, downregulating ANP32A restored synaptic plasticity and memory loss (Chai et al., 2017; Feng et al., 2017). The pathological role of ANP32A in Alzheimer’s disease contrasts with its protective role in cerebellar ataxia that we earlier demonstrated (Cornelis et al., 2018), suggesting that ANP32A has tissue-specific functions within the different brain regions. This supports the notion of the complexity of gene regulation in the human brain. In line with ANP32A’s pathological involvement in Alzheimer’s disease, ANP32A is reported to be increased in brain of Alzheimer’s disease patients and in disease mouse models (Tanimukai et al., 2005; Tsujio et al., 2005). In addition, overexpression of ANP32A in hippocampus induced memory impairments in mice (Chai et al., 2018). However, ANP32A’s pathological roles were not previously linked to a deficit in Wnt signaling, which has been extensively demonstrated to contribute to cognitive decline in Alzheimer’s disease (De Ferrari et al., 2014; Tapia-Rojas & Inestrosa, 2018). Thus, our study sheds light on the molecular mechanism underlying ANP32A link to Alzheimer’s disease, and supports that targeting ANP32A may prevent memory deficits by restoring the Wnt signaling balance, which is often lost in the ageing brain (Palomer et al., 2019).

This study has some limitations that are worth mentioning. *In vivo* experiments were performed with mice with a global deletion of the *Anp32a* gene as a conditional *Anp32a* is currently unavailable. Although unlikely, it may be possible that features reported here are not primarily caused by the absence of ANP32A in the particular explored tissue, but secondarily to other events occurring in other tissues. Thus, this study stimulates research in specific fields to corroborate the mechanisms reported here in a more tissue-specific manner. In addition, translation of preclinical interventions in mouse models, in particular of osteoarthritis, has been challenging. Many factors may play a role in this. Among these it is worth noting that the existence of multiple human osteoarthritis endophenotypes is not well represented in the post-traumatic DMM model, and that the impact of ageing on the chondrocyte’s identity and molecular program is challenging to mimic in mice.

In conclusion, our study identifies that ANP32A controls the transcriptional response of Wnt signaling in cartilage, heart and hippocampus, thereby suggesting ANP32A as a therapeutic target for diseases associated with dysregulation in this central signaling cascade. This insight, together with our previous report discovering ANP32A as regulator of the antioxidant defense, position ANP32A as a critical node regulating diverse key networks in the cell. As ANP32A expression is dysregulated in several diseases, further research should focus on the factors that control ANP32A expression. Our study also provides preclinical evidence for augmented therapeutic efficacy of a combinatorial treatment with Wnt inhibitors and antioxidants in osteoarthritis. Future studies could explore this combination in large animal models for osteoarthritis, and in other pathologies in which Wnt hyper-activation and oxidative stress simultaneously occur.

## Materials and Methods

### Study design

The objective of this study was to determine how ANP32A protects against the development of osteoarthritis by regulating pathways beyond oxidative stress and whether such mechanisms also affect other tissues and organs. Cartilage tissue and cells from patients without known osteoarthritis and genetically engineered mice were used in *ex vivo* and *in vivo* studies using unchallenged ageing mice and a model of joint disease, combined with *in vitro* assays. Mouse models are reported following the principles of the ARRIVE guidelines (https://www.nc3rs.org.uk/arrive-guidelines) (Supplementary Table 2). For all experiments, the largest possible sample size was used limited by low breeding of *Anp32a* deficient mice. For the *in vivo* osteoarthritis model a power analysis was based on our own historical data and the OARSI score as primary outcome (Cornelis et al., 2018). Based on previous experiments with the antioxidant intervention in *Anp32a*^-/-^ mice (Cornelis et al., 2018), we used G-Power software: a partial eta-squared of 0.3 results in an effect size of 0.65 predicting 80% power with 30 mice in total (8 per group). Analyses of other organs were exploratory without specific power calculation. Data from all animals are reported in the Figures – specific exclusion of 3 animals identified as extreme outliers in the statistical analysis of Figure 4I is described below and the data points from these animals are highlighted in the Figure. No animals were excluded in the experiments with the exception of one wild-type animal in the heart/body weight measurements due to clear physical abnormalities. Mice were randomly assigned to the experimental groups where applicable. Pathology analysis was performed in a blinded fashion.

### Patient materials

Human articular chondrocytes were isolated from the hips of patients undergoing total hip replacement surgery after informed consent. The University Hospitals Leuven Ethics Committee and Biobank Committee (Leuven, Belgium) approved the study (S56271). Under Belgian Law and UZ Leuven’s biobank policies, the hip joints are considered human biological residual material. Only age and sex of the patients are being shared between the surgeons and the investigators involved in this study. The material is fully anonymized without links to the medical file. Non-osteoarthritic articular chondrocytes were obtained from patients undergoing hip replacement for osteoporotic or malignancy-associated fractures. Upon arrival of the specimen, these non-osteoarthritis hips were macroscopically evaluated to exclude signs of osteoarthritis. For samples from the RAAK study, ethical approval was obtained from the medical ethics committee of LUMC (P08.239) (Leiden, The Netherlands) and informed consent was obtained from all participants (den Hollander et al., 2017; Y. F. M. Ramos et al., 2014). Herein, participant details can be found. Gene expression data based on RNAseq were obtained as described before (Cornelis et al., 2018; Y. F. Ramos et al., 2014).

### Mice

*Anp32a^-/-^* (*Anp32a^tm1Hzo^*) mice were a kind gift from Dr. P. Opal (Northwestern University Medical School, Chicago, USA) (Opal et al., 2004) and were backcrossed onto the C57Bl/6J background. In the experiments reported here, mice were between the 11^th^ and 21^st^ generation of backcrossing. Wild-type C57Bl/6J, purchased from Janvier (Le Genest St Isle, France), were used as controls. Mice with normal immune status were housed in groups of 4-5 mice in Static micro-insulator cage with Macrolon filter with bedding material (composed of spruce particles of approximately 2.5 - 3.5 mm, type Lignocel® BK 8/15), under conventional laboratory conditions (14h light – 10h dark; 23+/-2°C), with standard mouse chow food (Sniff, Soest, Germany) and water provided *ad libitum*. All studies were performed with the approval from the Ethics Committee for Animal Research (P114-2008, P198-2012, P159-2016; KU Leuven, Belgium) (Licence number LA1210189). Genotypes of animals were confirmed by polymerase chain reaction (PCR) (Cornelis et al., 2018).

### Mouse osteoarthritis models

For assessment of the joints after ageing, *Anp32a^-/-^* female mice and WT mice were sacrificed at the age of 12 months. In the destabilization of the medial meniscus (DMM) model, mild instability of the knee was induced by transection of the medial menisco-tibial ligament of the right knee of male 8-week old mice (Glasson et al., 2007). Knees were analyzed 12 weeks after surgery. These male mice received N-acetyl-cysteine (NAC) (Zambon S.A./N.V.) for 11 weeks, starting from the second week after the DMM surgery. NAC was added to the drinking water at a concentration of 1 mg/ml and mice were allowed to drink *ad libitum*. The solution was refreshed 3 times per week and protected from light. Wnt inhibitor XAV939 (XAV, 0.5 mg kg–1) (Selleckem) (Monteagudo et al., 2017) or vehicle was injected intra-articularly in the right knee of DMM-operated male mice starting from the second week after the surgery, with an interval of 10 days for a total of 7 times. PBS was injected in the left knee.

### Histology of the mouse joint

Mice were sacrificed, and knees were fixed in 2% formaldehyde overnight at 4°C; decalcified for 3 weeks in 0,5M EDTA pH 7.5; and embedded in paraffin. Hematoxylin-safranin-O staining and immunohistochemistry was performed on 5 µm thick sections. Pictures were taken using a Visitron Systems microscope (Leica Microsystems GmbH) using Spot32 software. Severity of disease was determined by histological scores throughout the knee (6 sections at 100 µm distance). Cartilage damage was assessed based on OARSI guidelines: depth of lesion (0-6) was scored on frontal knee sections (Glasson et al., 2010). Lesion grades represent the following features; 0: surface and cartilage morphology intact, 1: small fibrillations without loss of cartilage, 2: vertical clefts below superficial layer and some loss of surface lamina, 3-6: vertical clefts/erosions to the calcified cartilage extending 3: less than 25%, 4: 25-50%, 5: 50-75% and 6: more than 75%. Medial and lateral tibial and femoral cartilage was scored and the score represents the mean of the four quadrants. Scoring was done blinded to the group assignment. A scoring system for osteophytes was developed. The presence and size of osteophytes were quantified at the four quadrants. Scoring grades represent the following features; 0: no osteophytes; 0.5: possible small osteophyte; 1: definite small osteophyte, 2: medium osteophyte, 3: large and mature osteophyte (Cornelis et al., 2019). Medial and lateral tibia and femur were scored and the score represents the mean of the 4 quadrants. Scoring was done blinded to the group assignment.

### Histology of the mouse heart and hippocampus

20-week old male mice were sacrificed, hearts were weighed and fixed in 4% formaldehyde overnight at 4°C and embedded in paraffin. 16-week old male mice were sacrificed, and brains were fixed in 4% formaldehyde overnight at 4°C and embedded in paraffin. Immunohistochemistry was performed on 5 µm thick sections. Pictures were taken using a Visitron Systems microscope (Leica Microsystems GmbH) using Spot32 software. Immunohistochemistry was performed on 5 µm thick sagittal sections and picrosirius red staining on 5 µm thick coronal sections. In brief, sections were deparaffinized, incubated in picrosirius red for 1 hour, washed in acetic acid solution, washed in absolute alcohol and mounted. Quantification of the percentage of fibrosis was performed with color deconvolution plugin (Jacqui Ross, Auckland University) in ImageJ Software (NIH Image, National Institutes of Health, Bethesda, MD, USA). Quantification was performed on 3 different intersections for 3 to 5 different mice per condition.

### Immunohistochemistry

Immunohistochemistry was performed on paraffin embedded EDTA decalcified mouse knee sections, and on non-decalcified heart and brain sections. First, heat induced epitope retrieval was performed using a Citrate-EDTA buffer (pH 6.2 for TCF1 and pH 6.0 for AXIN2) for 10 min at 95°C. Then, sections were treated with 3% H_2_O_2_/methanol for 10 minutes to inactivate endogenous peroxidase activity. The sections were blocked in normal goat serum for 30 minutes and incubated overnight at 4°C with the primary antibodies against TCF1 (Ab96777, Abcam; 10 µg/ml for knee; 7.5 µg/ml for heart and brain) or AXIN2 (Ab32197, Abcam; 1.5 µg/ml for knee). After incubation with the primary antibody, 1:100 biotin-conjugated goat anti-rabbit IgG (Jackson Immunoresearch) was applied for 30 minutes. An amplification step using the Vectastain ABC kit (Vector Laboratories). And peroxidase activity was determined using DAB (Dako). Rabbit IgG (Santa Cruz Biotechnologies) was used as negative control.

### Quantitative PCR

Total RNA was isolated using a manual Trizol+phenol/chloroform (Invitrogen) extraction protocol or with the NucleoSpin RNA extraction kit (Macherey-Nagel, Filterservices) according to the manufacturer’s instructions. cDNA was synthetized using 1 µg total RNA with the RevertAid H Minus First Strand cDNA Synthesis Kit (Fermentas) according to the manufacturer’s recommendations. The relative expression level of transcripts was determined by real-time RT-PCR using cDNA as a template with Maxima SYBR Green qPCR Master Mix (Fermentas), following manufacturer’s instructions and gene-specific primers on a Corbett Rotor-gene qPCR machine (Qiagen). Relative quantification was obtained with the ΔΔct method using Hprt or 29s as internal control, unless otherwise specified. Primers used in quantitative PCR are listed in Supplementary Table 3.

### ATDC5 cartilage differentiation model

ATDC5 cells (Sigma) were cultured in growth medium (1:1 Dulbecco’s modified Eagle’s medium (DMEM): Ham’s F-12 mix, Gibco) containing 1% (vol/vol) antibiotic–antimycotic (Gibco), 5% fetal bovine serum (FBS) (Gibco), 10 µg/mL human transferrin (Sigma) and 30 nM sodium selenite (Sigma). Cells were maintained in a humidified atmosphere of 5% CO_2_ at 37°C. Stable ATDC5 cell lines were established using a GIPZ-shRNA construct directed against mouse *Anp32a* (Open Biosystems, now Dharmacon; clone ID V3LMM_502656); and a non-interfering GIPZ vector (Dharmacon) was used as a control. ATDC5 cells were transfected using Lipofectamine LTX (Thermo Fisher) and after 24h, selection with 3 µg/ml puromycin (Thermo Fisher) was initiated and continued for 14 days. Three different antibiotic-resistant clonal colonies of each condition were isolated and grown independently. Knock-down efficiency was assessed by quantitative RT-PCR.

Stably transfected ATDC5 cell lines were cultured as high-density micromasses (Wang et al., 2019). Cells were trypsinised, washed and resuspended at 2×10^7^ cells/mL in a chondrogenic medium containing DMEM-F12 enriched by 1% (vol/vol) antibiotic–antimycotic, 5% FBS, 5 µg/mL human transferrin and 1× ITS (Insulin, Transferrin, Selenite) premix (resulting in 10 µg/mL insulin, 5 µg/mL human transferrin and 30 nM sodium selenite) (Life Technologies). One droplet (10 µL) was carefully placed in the center of each well of a 24-well plate. Cells were allowed to adhere for 2 hours at 37°C, followed by addition of 500 µL of chondrogenic medium. Medium was renewed daily, and after 14 days of culture, cells were switched to α-MEM medium (Gibco) (with 1% (vol/vol) antibiotic–antimycotic, 5% FBS, 5 µg/mL human transferrin, 1× ITS premix) supplemented with 50 µg/ml ascorbic acid-2-phosphate and 7 mM β-glycerolphosphate to induce hypertrophic differentiation and mineralization of the extracellular matrix.

For the rescue experiments, micromassess were treated with N-acetyl-cysteine (NAC) (Sigma Aldrich) or XAV939 (Selleckchem) at the indicated concentrations.

### Micromass staining

ATDC5 micromasses were washed with PBS and fixed with 95% ice-cold methanol for 30 min at 4°C. After rinsing with water, micromasses were stained with either alcian blue (AB) (0.1% AB 8GX (Sigma)), safranin O (SO) (Klinipath), sirius red (SR) (0.1% Direct Red 80 (Sigma) in a saturated aqueous solution of picric acid) or alizarin red (AR) (1% AR (Sigma) in water pH 4.2), washed with water to remove unbound staining and air dried.

### Human Articular Chondrocyte Culture and transfection

Human articular chondrocytes were obtained from patients undergoing hip replacement for osteoporotic or malignancy-associated fractures. Cartilage was dissected from the joint explant surfaces and then rinsed with PBS. The tissue was cut into small pieces, using a sterile surgical blade. Cartilage explants were incubated with 2 mg/ml pronase solution (Roche) for 90 minutes at 37°C and digested overnight at 37°C in 1.5 mg/ml collagenase B solution (Roche) under continuous agitation. The preparation was filtered through a 70 µM strainer and cells were plated in culture flasks and cultured in a humidified atmosphere at 37°C, 5% CO_2_. Culture medium consisted of DMEM/F12 (Gibco), 10% fetal bovine serum (FBS) (Gibco), 1% (vol/vol) antibiotic/antimycotic (Gibco) and 1% L-glutamine (Gibco).

For small interfering RNA (siRNA) transfection, lipofectamin RNAiMAX (Invitrogen) was used as transfection reagent, together with siGENOME *ANP32A* siRNA (Dharmacon) or negative control siGENOME siRNA (Dharmacon), following the protocols provided by the manufacturer. Where indicated, cells were treated with 100 ng/ml recombinant WNT3A (R&D Systems).

### Luciferase assay

Primary human chondrocytes were plated into 96-well plates and after 24 h, cells were transfected with Super8X TOPFlash or Super8X FOPFlash (TOPFlash mutant control) (Addgene plasmids #12456 and #12457, respectively) for canonical Wnt signaling reporter using Lipofectamine LTX Reagent with PLUS Reagent (Life Technologies), according to the manufacturer’s protocol. After 24 h, cells were transfected with *ANP32A* or negative control siRNA, and 48 h later cells were treated with rWNT3A and collected 24 h later. Luciferase assay was performed using Promega Luciferase Assay system according to the manufacturer’s protocol. Briefly, cells were lysed with 400 µl per well 1 X lysis buffer (E1531) and underwent two freeze-thaw cycles to ensure complete cell lysis. Afterwards, 20 µl of cell lysate was transferred to opaque 96-well plates, and 50 µl of Luciferase Assay Reagent was dispensed into the plate. Luciferase level was measured using Plate-Reading Luminoskan Ascent (Thermo Scientific).

### Co-Immunoprecipitation

Co-Immunoprecipitation experiments were performed using the Pierce Co-Immunoprecipitation Kit (Thermo Scientific). Columns were conditioned following the manufacturer’s recommendations. Antibody binding to the column was performed using 75 mg of either a mock antibody (donkey anti-goat IgG) as a control or ANP32A antibody (Novus Biologicals, NBP1-97576, clone RJ1). After antibody immobilization, the columns were washed, and 100 mg of the lysate’s proteins were incubated overnight at 4°C under constant mixing. After three washings, retained proteins were eluted using 40 ml of Elution Buffer (Thermo Fisher), and stored at -20°C. Protein complexes were then detected by Western blotting.

### Immunofluorescence

Primary human chondrocytes were plated in 8-well glass chamber slides (Thermo) and treated with vehicle or recombinant WNT3A for 4 hours. After incubation, cells were fixed with 4% formaldehyde and permeabilized with 0.1% Triton-X-100. Rabbit polyclonal anti-ANP32A antibody (Abcam, ab5992, dilution 1:100) was added incubated overnight at 4°C. Alexa Flour 488-conjugated goat anti-rabbit secondary antibody (Abcam, ab150077, dilution 1:1000) was added and incubated for 1 h at room temperature. Nuclei were stained with Hoechst 33342 (Thermo, dilution 1:10000). The stainings were imaged by wide-field fluorescence microscopy using an Olympus IX83 inverted microscope (Olympus) equipped with a DP73 camera.

### ChIP Analysis

Chromatin immunoprecipitation assays were carried out using the Agarose ChIP kit from Thermo Scientific, according to the manufacturer’s guidelines. Briefly, cell samples were crosslinked by 1% Formaldehyde for 10 min, and the reaction was stopped by the addition of glycine to a 125 mM final concentration. The fixed cells were lysed in SDS buffer, and the chromatin was fragmented by microccocal nuclease digestion. The sheared chromatin was incubated with antibodies against ANP32A (Santa Cruz, sc-100767) and RNA polymerase II (Thermo Scientific, MA1-46093), and recovered by binding to protein A/G agarose. Eluted DNA fragments were used directly for qPCR. Primers used in quantitative PCR are listed in Supplementary Table 3.

### Cell lysis and western blotting

Cells were homogenized in IP Lysis/Wash buffer (Thermo Fisher) supplemented with 5% (vol/vol) Protease Mixture Inhibitor (Sigma) and 1mM phenylmethanesulfonyl (Sigma). After two homogenization cycles (7 s) with an ultrasonic cell disruptor (Microson; Misonix), total cell lysates were centrifuged 10 min at 18,000g, and supernatant containing proteins was collected. The protein concentration of the extracts was determined by Pierce BCA Protein Assay Kit (Thermo Scientific). Immunoblotting analyses were performed as described in previous studies (Cornelis et al., 2018; Monteagudo et al., 2017). Antibodies against AXIN1 (Millipore, 05-1579, clone A5, dilution 1:1000), Active β-catenin (Cell signaling, 8814S, cloneD13A1, dilution 1:1000), Total β-catenin (BD Transduction Laboratories, 610154, clone 14/Beta-Catenin, dilution 1:2000), ANP32A (Abcam, ab189110; dilution 1:1000), total H3 (Abcam, ab1791; dilution 1:10000) and Actin (Sigma, A2066; dilution 1:4000) were used following manufacturer’s instructions. The blotting signals were detected using the SuperSignalWest Femto Maximum Sensitivity Substrate system (Thermo Scientific).

### Bioinformatic analyses

Transcriptome analysis of articular cartilage from male *Anp32a*^-/-^ compared to WT mice was previously reported (Cornelis et al., 2018) (GEO nr: GSE108036). Differentially expressed genes (P<0.01) were analysed with Ingenuity Pathway Analysis software (Qiagen) applying the Upstream Regulator Analysis approach. Identified genes were then further evaluated for pathway enrichment using the PANTHER database version 11.1 (http://pantherdb.org) (Mi et al., 2017).

Tissue based gene expression profiles of *ANP32A* and *TCF1* were obtained from the Human Protein Atlas (http://www.proteinatlas.org) (Uhlén et al., 2015) and analysed with R-Studio (version 1.1.463). Expression data were log-transformed and scaled by subtraction of the variable mean and dividing the centered values by the variable standard-deviation). Data were organized by organ system and system with more than 3 organs or tissues represented were visualized using split plots.

### Statistics

Data analysis and graphical presentation were performed with R-Studio (version 1.1.463) and GraphPad Prism version 8. Power analysis was performed with G*Power (version 3.1.9.4). Data are presented as mean and SD or as individual data points, representing the mean of technical replicates as indicated in the figure legends. Raw data for the different experiments are available in Supplementary File 1. R code is available in Supplementary File 2. Gene expression data were log-transformed for statistical analysis. All test performed were two-tailed where applicable.

Distribution of the dependent variables was assessed by histogram inspection and boxplots. Univariate outliers and extreme outliers were identified as values 1.5 or 3 times the interquartile range below or above quartile 1 or quartile 3 respectively. Multivariate outliers were evaluated using Mahalanobis distance. Extreme outliers were removed for the multivariate analysis as indicated in figure legends. Univariate normality was assessed by Shapiro test and QQ plots.

For comparisons between two groups, Student’s t-test assuming equal variance was used. For multi-dependent variable analysis, MANOVA (multivariate analysis of variance) was used. Collinearity was evaluated in a correlation matrix. When correlation was ≥ 0.9 between dependent variables, the number of dependent variables was reduced accordingly. Homogeneity of co-variances and variances were tested by Box-M test and Levene test respectively. When Levene test was < 0.01, the dependent variable was removed from the model, as indicated in figure legends. Pillai’s trace, F-value and corresponding P-value are reported in figure legends.

Two-way ANOVA (analysis of variances) with type III squares was used to study interactions and main effects between independent categorical variables with or without within-subject factors as appropriate. Data are reported by F-values with degrees of freedom and exact p-values for main effects and interaction in the figure legends where applicable. The Greenhouse-Geisser sphericity correction was applied to within-subject factors violating the sphericity assumption. Homogeneity of variance was evaluated by residuals versus fit plot and by the Levene test. For comparisons between two groups, Student’s t-test assuming equal variance was used.

Expression data from the RAAK dataset were analyzed as published before (Cornelis et al., 2018), with differential gene expression being tested pairwise between preserved and lesioned samples by paired t-tests (preserved and damaged area per individual patient) followed by Benjamini-Hochberg correction to adjust for multiple testing. Uncorrected correlation analysis was performed for exploratory analysis of the relationship between Wnt target gene and ANP32A levels.

### Data availability

All data associated with this study are present in the paper or the Supplementary information. Microarray data were deposited to the Gene Expression Omnibus (www.ncbi.nlm.nih.gov/geo/) under accession no. GSE108036.

## Acknowledgments

We are grateful to L. Storms and A. Hens for their technical support in this study and for taking care of the animal facility management. We also thank M. Van Mechelen for her kind technical assistance. We are indebted to the UZ Leuven traumatology and orthopedic surgeons, and nursing staff (in particular M. Penninckx) for their efforts to provide cartilage samples for ex vivo and in vitro work, and to Prof. R.G.H.H Nelissen (LUMC) for collecting the samples of the Research Arthritis and Articular Cartilage (RAAK) study. We thank all study participants of the RAAK study. This work was supported by the Flanders Research Foundation (FWO-Vlaanderen), Excellence of Science (EOS), the Dutch Arthritis Association (DAA_10_1-402), the Dutch Scientific Research council NWO/ZonMW VICI scheme (nr. 91816631/528), and the Leiden University Medical Center.

## Author contributions

R.J.L. and S.M. planned the study and designed all the in vitro, ex vivo, and in vivo experiments. F.M.F.C. performed the animal experiments. S.M., X.W., A.R., T.P. and J.Q. performed the in vitro experiments. R.C.A. and I.M. performed the analysis of the RAAK study. R.J.L. is responsible for all the other statistical analyses. A.S provided essential materials. S.M., R.J.L. and F.M.F.C. wrote the manuscript.

## Conflict of interest

Leuven Research and Development, the technology transfer office of KU Leuven, has received consultancy and speaker fees and research grants on behalf of R.J.L. from Abbvie, Boehringer-Ingelheim, Celgene, Eli-Lilly, Galapagos, Janssen, MSD, Novartis, Pfizer, Samumed, and UCB. The other authors declare that they have no competing financial interests.

---

**Supplementary Figure 1.**
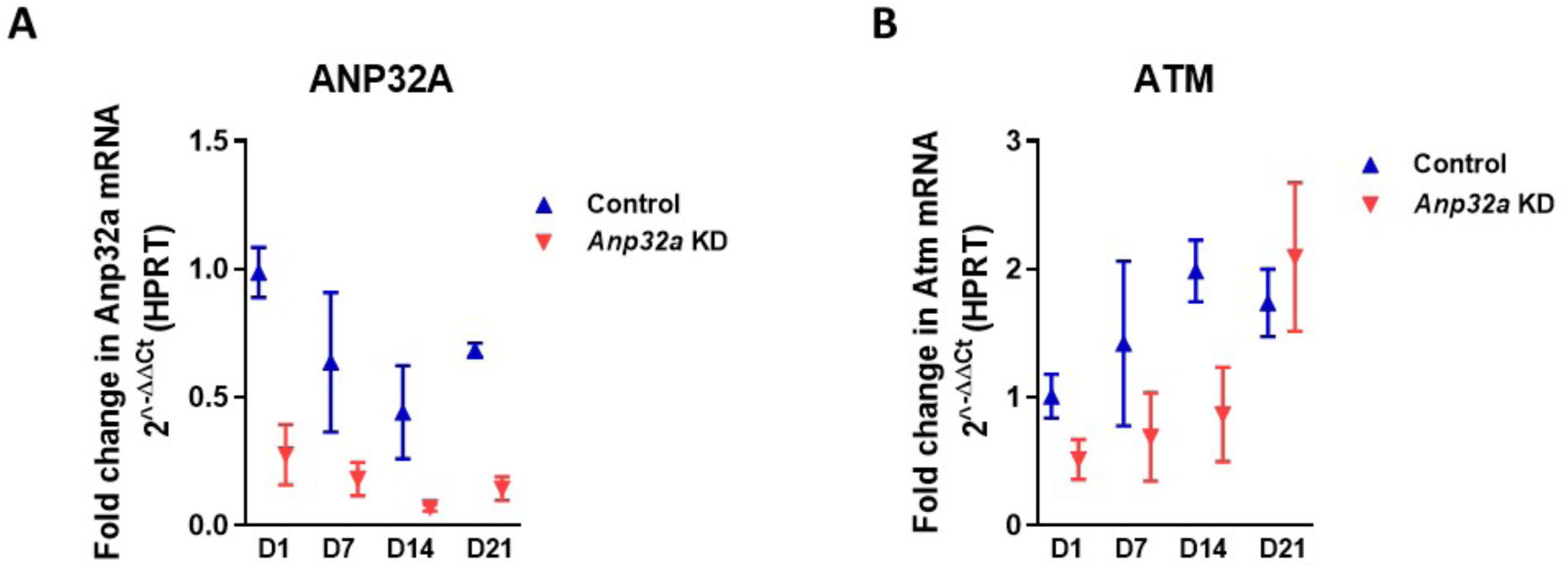
Gene expression analysis in ATDC5 micromasses. **(A, B)** Real-time PCR analysis of *Anp32a* (A) and *Atm* (B) in control and *Anp32a* knockdown (KD) ATDC5 cells. Error bars indicate mean ± SD of three technical replicates per condition [F_1,4_ = 69.892, *P* = 0.001 (*Anp32a*), F_1,4_ = 24.841 *P* = 0.009 (*Atm*), by 2-way ANOVA for control versus KD cells, mean ± SD of 3 technical replicates per condition].

**Supplementary Figure 2.**
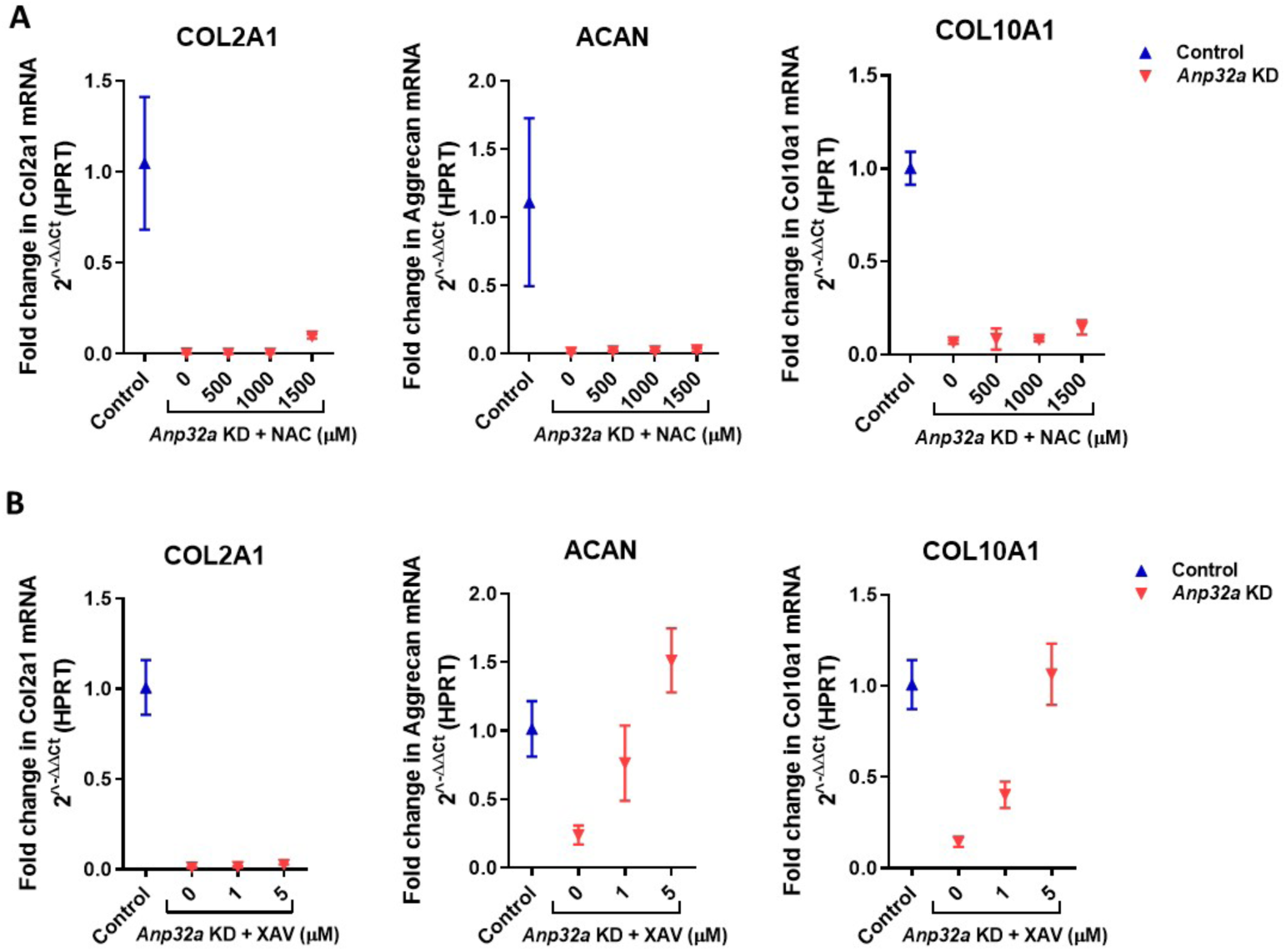
Rescue effects of antioxidant or Wnt inhibitor treatment in *Anp32a* knockdown ATDC5 micromasses. **(A, B)** Real-time PCR analysis of chondrogenic differentiation markers collagen 2 (*Col2a1*), aggrecan (*Acan*) and collagen 10 (*Col10a1*) at day 7 in control and *Anp32a* knockdown (KD) ATDC5 cells after treatment with antioxidant N-acetylcysteine (NAC) (A) or Wnt inhibitor XAV939 (XAV) (B) at the indicated concentrations. Error bars indicate mean ± SD of three technical replicates per condition.

**Supplementary Figure 3.**
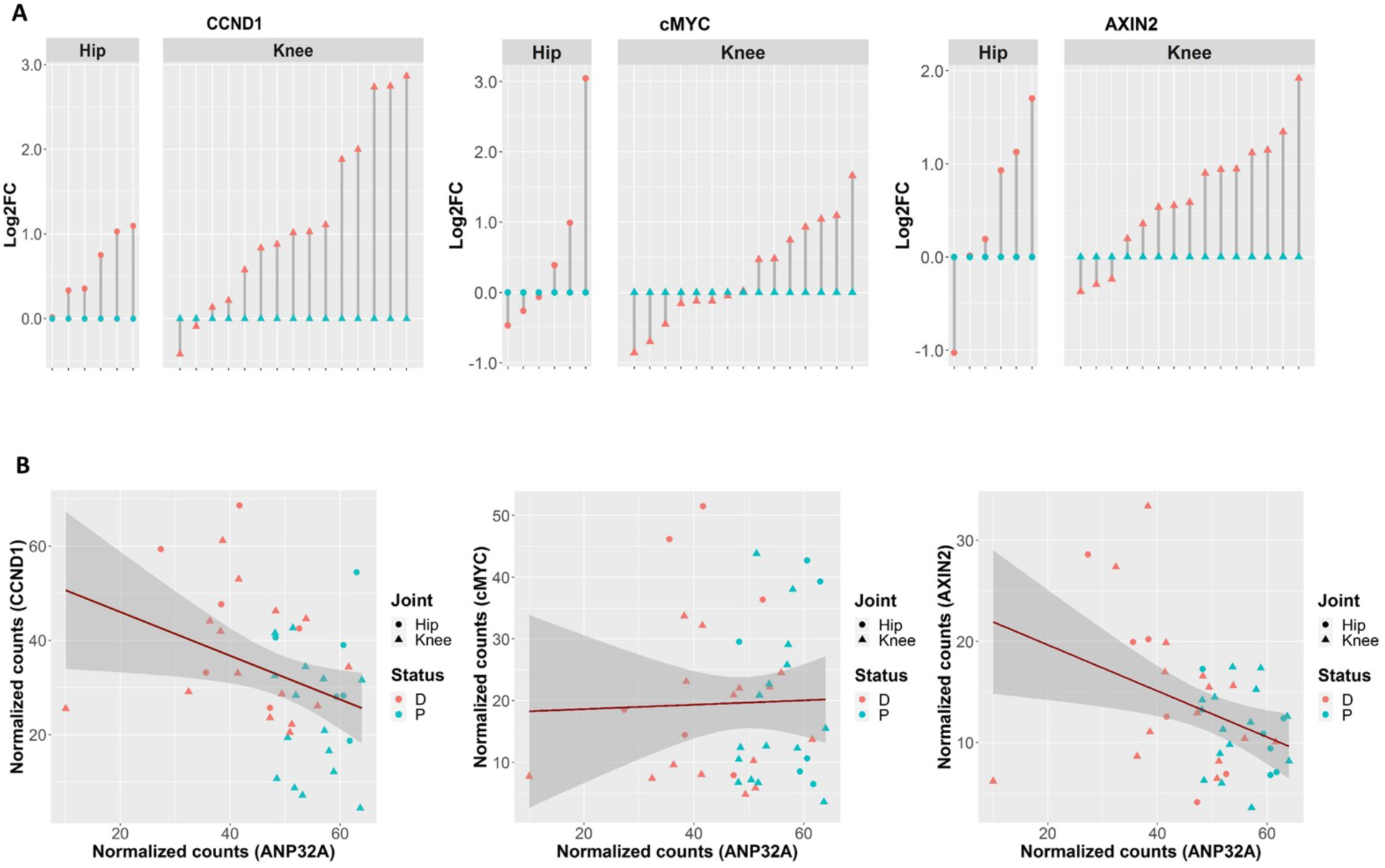
Expression of Wnt target genes in human osteoarthritis cartilage and correlation with *ANP32A*. **(A, B)** Expression of Wnt target genes *CCND1, CMYC and AXIN2* (A) and correlation with *ANP32A* expression (B) by RNA sequencing in paired preserved (P) and damaged (D) cartilage from hips (o) and knees (Δ) from osteoarthritis patients (log2-fold change (Log2FC) of damaged *vs.* preserved) (n=21, *P* < 0.0001 for *CCND1* and *AXIN2*, *P* =0.0809 for *CMYC* by Benjamini-Hochberg adjusted paired *t*-test (A), Pearson’s correlation R = -0.34 – *P =* 0.027, R = 0.03 – *P* = 0.853, R = -0.39 – *P* = 0.112 for *CCND1, CMYC and AXIN2* respectively (B)).

**Supplementary Table 1.** Body and heart weight, and ratio body/heart weight in *Anp32a*^-/-^ mice and WT controls.

|  |  | **Total body weight (g)** | **Heart weight (mg)** | **Ratio** |
| --- | --- | --- | --- | --- |
| **WT** | WT 1 | 26,6 | 137 | 5,150 |
|  | WT 2 | 29,5 | 141 | 4,780 |
|  | WT 3 | 28,9 | 152 | 5,260 |
|  | WT 4 | 26,8 | 141 | 5,261 |
|  | WT 5 | 26,9 | 146 | 5,428 |
|  | WT 6 | 24,0 | 152 | 6,333 |
|  | WT 7 | 24,0 | 115 | 4,792 |
|  | WT 8 | 31,4 | 138 | 4,395 |
|  | WT 9 | 27,6 | 136 | 4,928 |
|  | WT 10 | 31,8 | 140 | 4,403 |
|  | WT 11 | 31,0 | 136 | 4,387 |
|  | WT 12 | 31,4 | 129 | 4,108 |
|  | WT 13 | 26,2 | 117 | 4,466 |
|  | WT 14 | 24,2 | 123 | 5,083 |
|  | WT 15 | 24,1 | 127 | 5,270 |
| ***Anp32a*<sup>-/-</sup>** | KO 1 | 25,5 | 206 | 8,078 |
|  | KO 2 | 25,5 | 178 | 6,980 |
|  | KO 3 | 25,3 | 178 | 7,036 |
|  | KO 4 | 24,0 | 155 | 6,458 |
|  | KO 5 | 25,7 | 205 | 7,977 |
|  | KO 6 | 26,1 | 163 | 6,245 |
|  | KO 7 | 26,4 | 207 | 7,841 |
|  | KO 8 | 26,0 | 188 | 7,231 |
|  | KO 9 | 17,1 | 115 | 6,725 |
|  | KO 10 | 26,6 | 196 | 7,368 |
|  | KO 11 | 27,1 | 194 | 7,159 |
|  | KO 12 | 27,5 | 231 | 8,400 |
|  | KO 13 | 24,1 | 159 | 6,598 |
|  | KO 14 | 25,1 | 218 | 8,685 |
|  | KO 15 | 27,1 | 192 | 7,085 |
|  | KO 16 | 28,1 | 189 | 6,726 |
|  | KO 17 | 27,1 | 191 | 7,048 |
|  | KO 18 | 25,5 | 202 | 7,922 |
|  | KO 19 | 26,0 | 186 | 7,154 |
|  | KO 20 | 23,1 | 183 | 7,922 |
|  | KO 21 | 27,0 | 193 | 7,148 |
|  | KO 22 | 25,1 | 230 | 9,163 |
|  | KO 23 | 28,4 | 193 | 6,796 |
|  | KO 24 | 27,2 | 209 | 7,684 |
|  | KO 25 | 25,8 | 185 | 7,171 |
|  | KO 26 | 24,9 | 154 | 6,185 |
|  | KO 27 | 24,9 | 148 | 5,944 |
|  | KO 28 | 26,1 | 184 | 7,050 |
|  | KO 29 | 27,2 | 151 | 5,551 |
|  | KO 30 | 25,6 | 228 | 8,906 |
|  | KO 31 | 26,2 | 171 | 6,527 |
|  | KO 32 | 26,5 | 222 | 8,377 |
|  | KO 33 | 25,7 | 156 | 6,070 |
|  | KO 34 | 25,0 | 150 | 6,000 |
|  | KO 35 | 27,1 | 202 | 7,454 |

**Supplementary Table 2.** Animal experiments: Overview, setup, and analysis details. Abbreviations: AC, articular cartilage; qPCR, quantitative polymerase chain reaction; WT, wild-type; KO, knock-out; IHC, immunohistochemistry; DMM, destabilization of the medial meniscus; OA, osteoarthritis; NAC, N-acetyl-cysteine; XAV, XAV939.

| Experiment ID | Experiment details |
| --- | --- |
| 1. Tibia AC for qPCR | * 8-week old male C57Bl/6J and *Anp32a*<sup>-/-</sup> mice<br>* Total sample size: n=15; WT: n=8, KO: n=7<br>* Primary outcome: qPCR detection of mRNA expression: Fig.1f |
| 2. 8-week old healthy knee | * 8-week old male C57Bl/6J and *Anp32a*<sup>-/-</sup> mice<br>* Total sample size: n=6; WT: n=3, KO: n=3<br>* Primary outcome: IHC detection of protein expression: Fig.1g |
| 3. DMM OA model, untreated | * 20-week old male C57Bl/6J and *Anp32a*<sup>-/-</sup> mice (time of sacrifice - induction of model at 8 weeks)<br>* Total sample size: n=6; WT DMM: n=3, KO DMM: n=3<br>* Primary outcome: IHC detection of protein expression: Fig.1h |
| 4. Ageing OA model | * 12-month old female C57Bl/6J and *Anp32a*<sup>-/-</sup> mice<br>* Total sample size: n=6; WT: n=3, KO: n=3<br>* Primary outcome: IHC detection of protein expression: Fig.1i |
| 5. DMM OA model: dual therapy | * 20-week old male *Anp32a*<sup>-/-</sup> mice (time of sacrifice - induction of model at 8 weeks)<br>* Total sample size: n=32; Vehicle: n=8, NAC: n=8, XAV: n=8, NAC-XAV: n=8<br>* Primary outcome: OARSI grade: Fig.3c<br>* Primary outcome: Osteophyte grade: Fig.3f<br>* Secondary outcome: Histology of knees: Fig.3b and 3e<br>* Secondary outcome: IHC detection of protein expression in knees: Fig.3d and 3g<br>* Secondary outcome: qPCR detection of mRNA expression: Fig.4b, 4f<br>* Secondary outcome: IHC detection of protein expression in heart: Fig.4c<br>* Secondary outcome: Histology of heart: Fig.4d<br>* Secondary outcome: Heart weight: Fig.4e (KO + Vehicle: n=8), Supplementary Table 1 |
| 6. 20-week old *Anp32a*<sup>-/-</sup> mice | * 20-week old male *Anp32a*<sup>-/-</sup> mice<br>* Total sample size: n=3<br>* Primary outcome: Fibrosis staining and quantification heart: Fig.4g, 4h<br>* Secondary outcome: Heart weight: Fig.4e, Supplementary Table 1 |
| 7. 20-week old WT mice | * 20-week old male C57Bl/6J mice<br>* Total sample size: n=15<br>* Primary outcome: Histology of heart: Fig.4d<br>* Secondary outcome: Heart weight: Fig.4e, Supplementary Table 1<br>* Secondary outcome: IHC detection of protein expression in heart: Fig.4c<br>* Secondary outcome: Fibrosis staining and quantification heart: Fig.4g, 4h<br>* Secondary outcome: qPCR detection of mRNA expression: Fig.4b, 4f |
| 8. Brain for qPCR | * 8-week old male C57Bl/6J and *Anp32a*<sup>-/-</sup> mice<br>* Total sample size: n=18; WT: n=9, KO: n=9<br>* Primary outcome: qPCR detection of mRNA expression: Fig.4i |
| 9. Brain for IHC | * 16-week old male C57Bl/6J and *Anp32a*<sup>-/-</sup> mice<br>* Total sample size: n=6; WT: n=3, KO: n=3<br>* Primary outcome: IHC detection of protein expression: Fig.4j |

**Supplementary Table 3.**
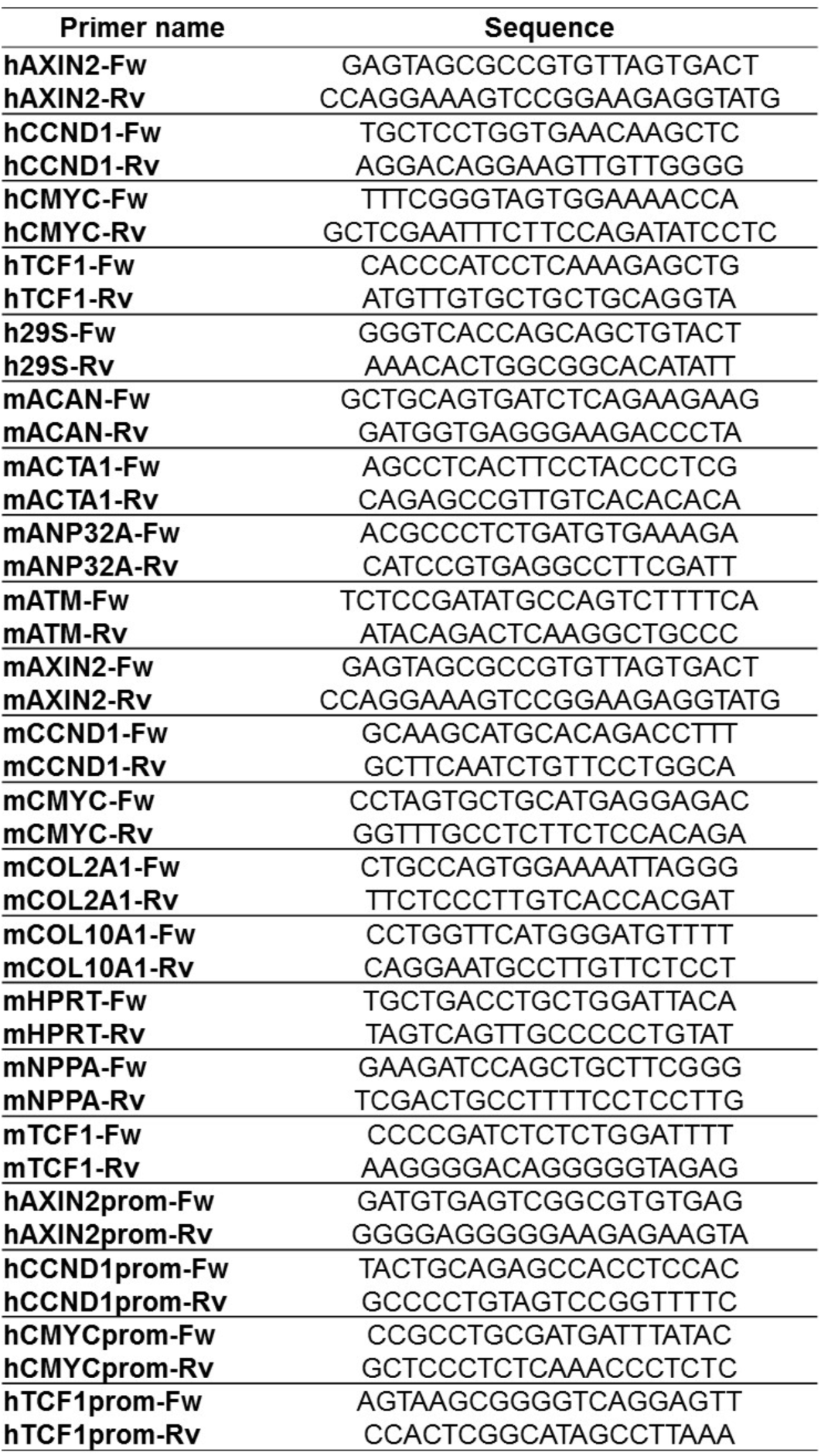
Primers used in qPCR analysis.

