## Supplementary file 2 for "ANP32A represses Wnt signaling across tissues tissues thereby protecting against joint and heart disease"

```r
# Packages required
library(car) # regression tools
library(rstatix) # pipe friendly data analysis
library(tidyverse) # facilitate data processing including ggplot2 graphics
library(ggpubr) # additional graph features
library(emmeans) # estimates and confidence intervals

# MANOVA script
# Analysis used for figure 1B - figure 1F - Figure 4B - Figure 4I
# Multiple dependent variables analysis of 4 Wnt target genes
# Four target genes of interest: TCF1, CCND1, cMYC, AXIN2
# Groups are wildtype or control vs ANP32 knockout or knockdown
# Outline of the procedure
# Log-transform gene expression data
# View histograms
# Test correlation matrix: if 0.9 > collinearity and remove - if #dep.variable = 2 analyse directly
# Test univariate normality
# Test multivariate normality
# Detect outliers
# Remove extreme outliers
# Test homogeneity of co-variance
# Test homogeneity of variance
# Perform MANOVA test

# t-test script
# Analysis used for figure 4E - figure 4F
# Groups are wildtype vs ANP32a knockout
# Outline of the procedure
# Log-transform gene expression data as per Vandesompele
# "https://blog.qbaseplus.com/seven-tips-for-bio-statistical-analysis-of-gene-expression-data"
# View histograms
# Test univariate normality
# Detect outliers
# Test homogeneity of variance
# Perform t-test

# 2-way ANOVA script with within subject effects (time points)
# Analysis used for figure 1C
# Groups are controls vs ANP32a knockdown
```

```
# Outline of the procedure
# Log-transform gene expression data as per Vandesompele
# "https://blog.qbaseplus.com/seven-tips-for-bio-statistical-analysis-of-gene-expression-
data"
# View histograms
# Test univariate normality
# Detect outliers
# Test homogeneity of variance
# Perform two-way ANOVA
# Geisser-Greehnouse sphericity correction automatically applied

# 2-way ANOVA scriptwith interaction between treatments
# Analysis used for figure 3C - 3F
# Groups are controls vs ANP32a-/- mice
# Outline of the procedure
# View histograms
# Perform lm regresson with interaction (= two-way anova)
# Use type III squares (unbalanced design)
# Test assumptions: residuals plot, QQ plot, levene test, shapiro test on residuals

# Figure 1B - file = figure1B.csv
# MANOVA test - multiple related dependent variables (wnt target genes)
target <- Figure1B
target$Group <- as.factor(target$Group) # set Group as factor
str(target) # view structure of the dataset
#transform
target2 <- target
target2$TCF1 <- log(target2$TCF1)
target2$CCND1 <- log(target2$CCND1)
target2$CMYC <- log(target2$CMYC)
target2$AXIN2 <- log(target2$AXIN2)
#view histograms and transformed Boxplot
hist(target2$TCF1)
hist(target2$CCND1)
hist(target2$CMYC)
hist(target2$AXIN2)
ggboxplot(target2, x = "Group", y = c("TCF1", "CCND1", "CMYC", "AXIN2"),
        merge = TRUE, palette = "jco")
# Assumptions testing
# identify outliers within dependent variables
target2 %>% group_by(Group) %>% identify_outliers(TCF1)
```

```
target2 %>% group_by(Group) %>% identify_outliers(CCND1)
target2 %>% group_by(Group) %>% identify_outliers(CMYC)
target2 %>% group_by(Group) %>% identify_outliers(AXIN2)
# identify multivariate outliers
target %>% group_by(Group) %>% mahalanobis_distance(-ID) %>% filter(is.outlier == TRUE)
%>% as.data.frame()
# univariate normality
target2 %>% group_by(Group) %>% shapiro_test(TCF1, CCND1, CMYC, AXIN2) %>%
arrange(variable)
# QQ plot of TCF1
ggqqplot(target2, "TCF1", facet.by = "Group",
        ylab = "TCF1", ggtheme = theme_bw())
ggqqplot(target2, "CCND1", facet.by = "Group",
        ylab = "CCND1", ggtheme = theme_bw())
ggqqplot(target2, "CMYC", facet.by = "Group",
        ylab = "CMYC", ggtheme = theme_bw())
ggqqplot(target2, "AXIN2", facet.by = "Group",
        ylab = "AXIN2", ggtheme = theme_bw())
# correlation matrix (must be < 0.9) cor_mat from Rstatic
target2 %>% cor_mat(TCF1, CCND1, CMYC, AXIN2) # if 0.9 remove variable and redo
# TCF1 - CCNDA 0.9 - TCF1 - AXIN2 0.9
# to reduce dimensions
target2 <- target2[,-c(4,6)]
head(target2)
#homogeneity of co-variances
box_m(target2[, c( "TCF1", "CMYC")], target2$Group)
# homogeneity of variance
target2 %>%
  gather(key = "variable", value = "value", TCF1, CMYC) %>%
  group_by(variable) %>%
  levene_test(value ~ Group)
# Run MANOVA
wnttarget <- manova(cbind(TCF1, CMYC) ~ Group, target2)
# Obtain results
wnttarget
summary(wnttarget, "Pillai")
summary.aov(wnttarget)

#Figure 1C - files = figure1Ccol2a1.csv - Figure1Cacan.csv - Figure1Ccol10a1.csv
COL2 <- Figure1Ccol2a1
ACAN <- Figure1Cacan
```

```
COL10 <- Figure1Ccol10a1
# Gather columns different times into long format
COL2long <- gather(COL2, key = "time", value = "genex", d1, d7, d14, d21)
ACANlong <- gather(ACAN, key = "time", value = "genex", d1, d7, d14, d21)
COL10long <- gather(COL10, key = "time", value = "genex", d1, d7, d14, d21)
# Set Factors
COL2long$time <- factor(COL2long$time)
ACANlong$time <- factor(ACANlong$time)
COL10long$time <- factor(COL10long$time)
# Check dataset
hist(COL2long$genex)
hist(ACANlong$genex)
hist(COL10long$genex)
# make log-transformed dependent variables
COL2xlong <- COL2long
ACANxlong <- ACANlong
COL10xlong <- COL10long
COL2xlong$genex <- log(COL2xlong$genex)
ACANxlong$genex <- log(ACANxlong$genex)
COL10xlong$genex <- log(COL10xlong$genex)
# Check dataset
hist(COL2xlong$genex)
hist(ACANxlong$genex)
hist(COL10xlong$genex)
# identify outliers
COL2xlong %>% group_by(Group, time) %>% identify_outliers(genex)
ACANxlong %>% group_by(Group, time) %>% identify_outliers(genex)
COL10xlong %>% group_by(Group, time) %>% identify_outliers(genex)
# normal distruction
COL2xlong %>% group_by(Group, time) %>% shapiro_test(genex)
ACANxlong %>% group_by(Group, time) %>%  shapiro_test(genex)
COL10xlong %>% group_by(Group, time) %>% shapiro_test(genex)
# normal distruction
ggqqplot(COL2xlong, "genex", ggtheme = theme_bw()) +
  facet_grid(time ~ Group, labeller = "label_both")
ggqqplot(ACANxlong, "genex", ggtheme = theme_bw()) +
  facet_grid(time ~ Group, labeller = "label_both")
ggqqplot(COL10xlong, "genex", ggtheme = theme_bw()) +
  facet_grid(time ~ Group, labeller = "label_both")
# RUN ANOVAS - balanced design (type I, II, III squares identical)
COL2.aov <- anova_test(data = COL2xlong, dv = genex, wid = ID, between=Group,
```

```
                   within = time, type=2, detailed=FALSE)
ACAN.aov <- anova_test(data = ACANxlong, dv = genex, wid = ID, between=Group,
                   within = time, type=2, detailed=FALSE)
COL10.aov <- anova_test(data = COL10xlong, dv = genex, wid = ID, between=Group,
                    within = time, type=2, detailed=FALSE)
# Results
get_anova_table(COL2.aov, correction = c("auto"))
get_anova_table(ACAN.aov, correction = c("auto"))
get_anova_table(COL10.aov, correction = c("auto"))

# Figure 1F - file = figure1F.csv
# MANOVA test - multiple related dependent variables (wnt target genes)
target <- Figure1F
target$Group <- as.factor(target$Group) # set Group as factor
str(target) # view structure of the dataset
#transform
target2 <- target
target2$TCF1 <- log(target2$TCF1)
target2$CCND1 <- log(target2$CCND1)
target2$CMYC <- log(target2$CMYC)
target2$AXIN2 <- log(target2$AXIN2)
#view histograms and transformed Boxplot
hist(target2$TCF1)
hist(target2$CCND1)
hist(target2$CMYC)
hist(target2$AXIN2)
ggboxplot(target2, x = "Group", y = c("TCF1", "CCND1", "CMYC", "AXIN2"),
         merge = TRUE, palette = "jco")
# Assumptions testing
# identify outliers within dependent variables
target2 %>% group_by(Group) %>% identify_outliers(TCF1)
target2 %>% group_by(Group) %>% identify_outliers(CCND1)
target2 %>% group_by(Group) %>% identify_outliers(CMYC)
target2 %>% group_by(Group) %>% identify_outliers(AXIN2)
# identify multivariate outliers
target %>% group_by(Group) %>% mahalanobis_distance(-ID) %>% filter(is.outlier == TRUE)
%>% as.data.frame()
# univariate normality
target2 %>% group_by(Group) %>% shapiro_test(TCF1, CCND1, CMYC, AXIN2) %>%
arrange(variable)
# QQ plot of TCF1
```

```
ggqqplot(target2, "TCF1", facet.by = "Group",
        ylab = "TCF1", ggtheme = theme_bw())
ggqqplot(target2, "CCND1", facet.by = "Group",
        ylab = "CCND1", ggtheme = theme_bw())
ggqqplot(target2, "CMYC", facet.by = "Group",
        ylab = "CMYC", ggtheme = theme_bw())
ggqqplot(target2, "AXIN2", facet.by = "Group",
        ylab = "AXIN2", ggtheme = theme_bw())
# correlation matrix (must be < 0.9) cor_mat from Rstatic
target2 %>% cor_mat(TCF1, CCND1, CMYC, AXIN2) # if 0.9 remove variable and redo
# TCF1 - CMYC 0.9
# to reduce dimensions
target2 <- target2[,-c(5)]
head(target2)
# homogeneity of co-variances
box_m(target2[, c( "TCF1", "CCND1", "AXIN2")], target2$Group)
# homogeneity of variance
target2 %>%
  gather(key = "variable", value = "value", TCF1, CCND1,AXIN2) %>%
  group_by(variable) %>%
  levene_test(value ~ Group)
# Run MANOVA
wnttarget <- manova(cbind(TCF1, CCND1,  AXIN2) ~ Group, target2)
# Obtain results
wnttarget
summary(wnttarget, "Pillai")
summary.aov(wnttarget)

# Figure 3C - file: Figure3C.csv
# OARSI score average of 4 quadrants by 2-way ANOVA with interaction
oarsi <- Figure3C
# Set variables NAC - XAV as factors
oarsi$XAV <- as.factor(oarsi$XAV)
oarsi$NAC <- as.factor(oarsi$NAC)
oarsi$treatment <- as.factor(oarsi$treatment)
str(oarsi)
# visualize dependent variable
hist(oarsi$meanfour)
# Box plots with two factor variables
boxplot(meanfour ~ NAC * XAV, data=oarsi, frame = FALSE,
       col = c("#00AFBB", "#E7B800"), ylab="OARSI score")
```

```
# Two-way interaction plots
interaction.plot(x.factor = oarsi$NAC, trace.factor = oarsi$XAV,
            response = oarsi$meanfour, fun = mean,
            type = "b", legend = TRUE,
            xlab = "NAC", ylab="OARSI score",
            pch=c(1,19), col = c("#00AFBB", "#E7B800"))
# run the models - lm function
meanfour.aov <- lm(meanfour ~ XAV*NAC, data=oarsi) # with interaction
summary(meanfour.aov) # evaluate overall strength of linear model
# contrasts
options(contrasts = c("contr.sum", "contr.poly")) # unbalanced design > adapt contrasts for
Anova function
Anova(meanfour.aov, type = "II") # give data summary with Anova function
# determine estimates and confidence intervals with "emmeans"
confint(emmeans(meanfour.aov, pairwise ~ "NAC"))
# Check assumptions
plot(meanfour.aov, 1) # homogeneity of variances
plot(meanfour.aov, 2) # qq plot - normal distribution
leveneTest(meanfour ~ XAV*NAC, data = oarsi) # Levene test
aovmf_residuals <- residuals(object = meanfour.aov) # Extract residuals
shapiro.test(x = aovmf_residuals ) # Run Shapiro-Wilk test
outlierTest(meanfour.aov) # tests outliers

# Figure 3F - file: Figure3F.csv
# osteophytes score of in medial compartment by 2-way ANOVA with interaction
osteo <- Figure3F
# Set variables NAC - XAV as factors
osteo$XAV <- as.factor(osteo$XAV)
osteo$NAC <- as.factor(osteo$NAC)
osteo$treatment <- as.factor(osteo$treatment)
str(osteo)
# visualize dependent variable
hist(osteo$osavm)
# Box plots with two factor variables
boxplot(osavm ~ NAC * XAV, data=osteo, frame = FALSE,
     col = c("#00AFBB", "#E7B800"), ylab="Osteophyte score")
# Two-way interaction plots
interaction.plot(x.factor = osteo$NAC, trace.factor = osteo$XAV,
            response = osteo$osavm, fun = mean,
            type = "b", legend = TRUE,
            xlab = "NAC", ylab="Osteophyte score",
```

```
pch=c(1,19), col = c("#00AFBB", "#E7B800"))
# run the models - lm function
osteo.aov <- lm(osavm ~ XAV*NAC, data=osteo) # with interaction
summary(osteo.aov) # evaluate overall strength of linear model
# contrasts
options(contrasts = c("contr.sum", "contr.poly")) # unbalanced design > adapt contrasts for Anova function
Anova(osteo.aov, type = "III") # give data summary with Anova function
# determine estimates and confidence intervals with "emmeans"
confint(emmeans(osteo.aov, pairwise ~ "NAC"))
# Check assumptions
plot(osteo.aov, 1) # homogeneity of variances
plot(osteo.aov, 2) # qq plot - normal distribution
leveneTest(osavm ~ XAV*NAC, data = osteo) # Levene test
aovmf_residuals <- residuals(object = osteo.aov) # Extract residuals
shapiro.test(x = aovmf_residuals ) # Run Shapiro-Wilk test
outlierTest(osteo.aov) # tests outliers

#Figure 4A
mydata <- Figure4A
head(mydata)
# define variables
mydata$Group <- as.factor(mydata$Group)
mydata$ID <- as.factor(mydata$ID)
mydata$Group <- as.factor(mydata$Group)
summary(mydata)
# show distributions
hist(mydata$ANP32A)
hist(mydata$TCF1)
# transform
mydata$logANP <- log(mydata$ANP32A)
mydata$logTCF <- log(mydata$TCF1)
# after transformation
hist(mydata$logANP)
hist(mydata$logTCF)
# scale the data
mydata$logANP2 <- scale(mydata$logANP, center = TRUE, scale = TRUE)
mydata$logTCF2 <- scale(mydata$logTCF, center = TRUE, scale = TRUE)
# visualise the data
(colour_plot <- ggplot(mydata, aes(x = logANP2, y = logTCF2)) +geom_point(size = 2) +
theme_classic2() +
```

```
   theme(legend.position = "none"))
# identify the strong outliers - no extreme outliers found
mydata %>% identify_outliers(logANP2)
mydata %>% identify_outliers(logTCF2)
mydata2 <- mydata
# make graph
(colour_plot <- ggplot(mydata2, aes(x = logANP2, y = logTCF2, colour=Group))
  + geom_point(size = 3) + theme_classic2() + xlab("log(ANP32a) Z-score") + ylab("log(TCF1)
Z-score") +
   theme(legend.position = "none") + coord_cartesian(xlim = c(-2.5, 2.5), ylim = c(-2.5, 2.5)))
# provide split graphs with groups
(split_plot <- ggplot(aes(logANP2, logTCF2, colour=Group), data = mydata) +
   geom_point() + facet_wrap(~ Group) + theme(legend.position = "none") +
xlab("log(ANP32A) Z-score") + ylab("log(TCF1) Z-score")
  + geom_hline(yintercept=0, size=0.3, linetype="dashed") + geom_vline(xintercept=0,
size=0.3, linetype="dashed") + coord_cartesian(xlim = c(-2.5, 2.5), ylim = c(-2.5, 2.5)))
# remove small groups
mydata2 <- mydata [-c(13,18, 30, 31, 42, 43, 47, 48),]
# update split graph
(split_plot <- ggplot(aes(logANP2, logTCF2, colour=Group), data = mydata2) +
   geom_point(size=3) + facet_wrap(~ Group, nrow=1) + theme(legend.position = "none") +
theme(aspect.ratio = 1) + theme(text = element_text(size = 11)) + xlab("log(ANP32A)
scaled") + ylab("log(TCF1) scaled")
  + geom_hline(yintercept=0, size=0.5, linetype="dashed") + geom_vline(xintercept=0,
size=0.5, linetype="dashed") + coord_cartesian(xlim = c(-2.5, 2.5), ylim = c(-2.5, 2.5)))

# Figure 4B - file = figure4B.csv
# MANOVA test - multiple related dependent variables (wnt target genes)
target <- Figure4B
target$Group <- as.factor(target$Group) # set Group as factor
str(target) # view structure of the dataset
#transform
target2 <- target
target2$TCF1 <- log(target2$TCF1)
target2$CCND1 <- log(target2$CCND1)
target2$CMYC <- log(target2$CMYC)
target2$AXIN2 <- log(target2$AXIN2)
#view histograms and transformed Boxplot
hist(target2$TCF1)
hist(target2$CCND1)
hist(target2$CMYC)
```

```
hist(target2$AXIN2)
ggboxplot(target2, x = "Group", y = c("TCF1", "CCND1", "CMYC", "AXIN2"),
          merge = TRUE, palette = "jco")
# Assumptions testing
# identify outliers within dependent variables
target2 %>% group_by(Group) %>% identify_outliers(TCF1)
target2 %>% group_by(Group) %>% identify_outliers(CCND1)
target2 %>% group_by(Group) %>% identify_outliers(CMYC)
target2 %>% group_by(Group) %>% identify_outliers(AXIN2)
# identify multivariate outliers
target %>% group_by(Group) %>% mahalanobis_distance(-ID) %>% filter(is.outlier == TRUE)
%>% as.data.frame()
# univariate normality
target2 %>% group_by(Group) %>% shapiro_test(TCF1, CCND1, CMYC, AXIN2) %>%
arrange(variable)
# QQ plot of TCF1
ggqqplot(target2, "TCF1", facet.by = "Group",
         ylab = "TCF1", ggtheme = theme_bw())
ggqqplot(target2, "CCND1", facet.by = "Group",
         ylab = "CCND1", ggtheme = theme_bw())
ggqqplot(target2, "CMYC", facet.by = "Group",
         ylab = "CMYC", ggtheme = theme_bw())
ggqqplot(target2, "AXIN2", facet.by = "Group",
         ylab = "AXIN2", ggtheme = theme_bw())
# correlation matrix (must be < 0.9) cor_mat from Rstatic
target2 %>% cor_mat(TCF1, CCND1, CMYC, AXIN2) # if 0.9 remove variable and redo
# homogeneity of co-variances
box_m(target2[, c( "TCF1", "CCND1", "CMYC", "AXIN2")], target2$Group)
# homogeneity of variance
target2 %>%
  gather(key = "variable", value = "value", TCF1, CCND1, CMYC, AXIN2) %>%
  group_by(variable) %>%
  levene_test(value ~ Group)
# Axin2 does not show homogeneity of variance
target2 <- target2[,-c(6)]
head(target2)
#homogeneity of co-variances
# homogeneity of co-variances
box_m(target2[, c( "TCF1", "CCND1", "CMYC")], target2$Group)
# homogeneity of variance
target2 %>%
```

```
  gather(key = "variable", value = "value", TCF1, CCND1, CMYC) %>%
  group_by(variable) %>%
  levene_test(value ~ Group)
# Run MANOVA
wnttarget <- manova(cbind(TCF1, CCND1,  CMYC) ~ Group, target2)
# Obtain results
wnttarget
summary(wnttarget, "Pillai")
summary.aov(wnttarget)

# Figure 4E - t-tests
mydata <- Figure4E
mydata$Group <- as.factor(mydata$Group) # set Group as factor
str(mydata) # view structure of the dataset
# view histograms and transformed boxplot
hist(mydata$Heart)
ggboxplot(mydata, x = "Group", y = "Heart",
          merge = TRUE, palette = "jco")
# Assumptions testing
# identify outliers within dependent variables
mydata %>% group_by(Group) %>% identify_outliers(Heart)
# univariate normality
mydata %>% group_by(Group) %>% shapiro_test(Heart) %>% arrange(variable)
# QQ plot of Heart
ggqqplot(mydata, "Heart", facet.by = "Group",
         ylab = "Heart", ggtheme = theme_bw())
# homogeneity of variance
mydata %>%
  gather(key = "variable", value = "value", Heart) %>%
  group_by(variable) %>%
  levene_test(value ~ Group)
# perform t-test with equal variance
t.test(Heart ~ Group, data = mydata, var.equal = TRUE)

# Figure 4F - t-tests
mydata <- Figure4F
mydata$Group <- as.factor(mydata$Group) # set Group as factor
str(mydata) # view structure of the dataset
mydata2 <- mydata
mydata2$NPPA <- log(mydata2$NPPA)
mydata2$ACTA1 <- log(mydata2$ACTA1)
```

```
# view histograms and transformed boxplot
hist(mydata2$NPPA)
hist(mydata2$ACTA1)
ggboxplot(mydata2, x = "Group", y = c("NPPA", "ACTA1"),
          merge = TRUE, palette = "jco")
# Assumptions testing
# identify outliers within dependent variables
mydata2 %>% group_by(Group) %>% identify_outliers(NPPA)
mydata2 %>% group_by(Group) %>% identify_outliers(ACTA1)
# univariate normality
mydata2 %>% group_by(Group) %>% shapiro_test(NPPA, ACTA1) %>% arrange(variable)
# QQ plot of NPPA
ggqqplot(mydata2, "NPPA", facet.by = "Group",
         ylab = "NPPA", ggtheme = theme_bw())
ggqqplot(mydata2, "ACTA1", facet.by = "Group",
         ylab = "ACTA1", ggtheme = theme_bw())
# correlation matrix (must be < 0.9) cor_mat from Rstatic
mydata2 %>% cor_test(NPPA, ACTA1) # if 0.9 remove variable and redo
# homogeneity of co-variances
box_m(mydata2[, c( "NPPA", "ACTA1")], mydata2$Group)
# homogeneity of variance
mydata2 %>%
  gather(key = "variable", value = "value", NPPA, ACTA1) %>%
  group_by(variable) %>%
  levene_test(value ~ Group)
# perform t-test with equal variance
t.test(NPPA ~ Group, data = mydata2, var.equal = TRUE)
t.test(ACTA1 ~ Group, data = mydata2, var.equal = TRUE)

# Figure 4H - t-tests
mydata <- Figure4H
mydata$Group <- as.factor(mydata$Group) # set Group as factor
str(mydata) # view structure of the dataset
# view histograms and transformed boxplot
hist(mydata$Fibrosis)
ggboxplot(mydata, x = "Group", y = "Fibrosis",
          merge = TRUE, palette = "jco")
# Assumptions testing
# identify outliers within dependent variables
mydata %>% group_by(Group) %>% identify_outliers(Fibrosis)
# univariate normality
```

```
mydata %>% group_by(Group) %>% shapiro_test(Fibrosis) %>% arrange(variable)
# QQ plot of Heart
ggqqplot(mydata, "Fibrosis", facet.by = "Group",
        ylab = "Fibrosis", ggtheme = theme_bw())
# homogeneity of variance
mydata %>%
  gather(key = "variable", value = "value", Fibrosis) %>%
  group_by(variable) %>%
  levene_test(value ~ Group)
# perform t-test with equal variance
t.test(Fibrosis ~ Group, data = mydata, var.equal = TRUE)

# Figure 4I - file = figure4I.csv
# MANOVA test - multiple related dependent variables (wnt target genes)
target <- Figure4I
target$Group <- as.factor(target$Group) # set Group as factor
str(target) # view structure of the dataset
#transform
target2 <- target
target2$TCF1 <- log(target2$TCF1)
target2$CCND1 <- log(target2$CCND1)
target2$CMYC <- log(target2$CMYC)
target2$AXIN2 <- log(target2$AXIN2)
#view histograms and transformed Boxplot
hist(target2$TCF1)
hist(target2$CCND1)
hist(target2$CMYC)
hist(target2$AXIN2)
ggboxplot(target2, x = "Group", y = c("TCF1", "CCND1", "CMYC", "AXIN2"),
         merge = TRUE, palette = "jco")
# Assumptions testing
# identify outliers within dependent variables
target2 %>% group_by(Group) %>% identify_outliers(TCF1)
target2 %>% group_by(Group) %>% identify_outliers(CCND1)
target2 %>% group_by(Group) %>% identify_outliers(CMYC)
target2 %>% group_by(Group) %>% identify_outliers(AXIN2)
# identify multivariate outliers
target %>% group_by(Group) %>% mahalanobis_distance(-ID) %>% filter(is.outlier == TRUE)
%>% as.data.frame()
# 3 extreme outliers to be removed ID 1 - 4 - 13
target2 <- target2[-c(1,4,16),]
```

```r
head(target2)
# univariate normality
target2 %>% group_by(Group) %>% shapiro_test(TCF1, CCND1, CMYC, AXIN2) %>%
arrange(variable)
# QQ plot of TCF1
ggqqplot(target2, "TCF1", facet.by = "Group",
        ylab = "TCF1", ggtheme = theme_bw())
ggqqplot(target2, "CCND1", facet.by = "Group",
        ylab = "CCND1", ggtheme = theme_bw())
ggqqplot(target2, "CMYC", facet.by = "Group",
        ylab = "CMYC", ggtheme = theme_bw())
ggqqplot(target2, "AXIN2", facet.by = "Group",
        ylab = "AXIN2", ggtheme = theme_bw())
# correlation matrix (must be < 0.9) cor_mat from Rstatic
target2 %>% cor_mat(TCF1, CCND1, CMYC, AXIN2) # if 0.9 remove variable and redo
# homogeneity of co-variances
box_m(target2[, c( "TCF1", "CCND1", "CMYC", "AXIN2")], target2$Group)
# homogeneity of variance
target2 %>%
  gather(key = "variable", value = "value", TCF1, CCND1, CMYC, AXIN2) %>%
  group_by(variable) %>%
  levene_test(value ~ Group)
# Run MANOVA
wnttarget <- manova(cbind(TCF1, CCND1, CMYC, AXIN2) ~ Group, target2)
# Obtain results
wnttarget
summary(wnttarget, "Pillai")
summary.aov(wnttarget)

#Figure S1 - files = FigureS1A.csv - FigureS1B.csv
ANP <- FigureS1A
ATM <- Figure1SB
# Gather columns different times into long format
ANPlong <- gather(ANP, key = "time", value = "genex", d1, d7, d14, d21)
ATMlong <- gather(ATM, key = "time", value = "genex", d1, d7, d14, d21)
# Set Factors
ANPlong$time <- factor(ANPlong$time)
ATMlong$time <- factor(ATMlong$time)
# Check dataset
hist(ANPlong$genex)
hist(ATMlong$genex)
```

```
# make log-transformed dependent variables
ANPxlong <- ANPlong
ATMxlong <- ATMlong
ANPxlong$genex <- log(ANPxlong$genex)
ATMxlong$genex <- log(ATMxlong$genex)
# Check dataset
hist(ANPxlong$genex)
hist(ATMxlong$genex)
# identify outliers
ANPxlong %>% group_by(Group, time) %>% identify_outliers(genex)
ATMxlong %>% group_by(Group, time) %>% identify_outliers(genex)
# normal distruction
ANPxlong %>% group_by(Group, time) %>% shapiro_test(genex)
ATMxlong %>% group_by(Group, time) %>%  shapiro_test(genex)
# normal distruction
ggqqplot(ANPxlong, "genex", ggtheme = theme_bw()) +
  facet_grid(time ~ Group, labeller = "label_both")
ggqqplot(ATMxlong, "genex", ggtheme = theme_bw()) +
  facet_grid(time ~ Group, labeller = "label_both")
# RUN ANOVAS - balanced design (type I, II, III squares identical)
ANP.aov <- anova_test(data = ANPxlong, dv = genex, wid = ID, between=Group,
                    within = time, type=2, detailed=FALSE)
ATM.aov <- anova_test(data = ATMxlong, dv = genex, wid = ID, between=Group,
                    within = time, type=2, detailed=FALSE)
# Results
get_anova_table(ANP.aov, correction = c("auto"))
get_anova_table(ATM.aov, correction = c("auto"))
```
